# Biocontrol of bacterial wilt disease using plant-associated bacterial communities in tomato

**DOI:** 10.1101/2024.09.14.613082

**Authors:** Eriko Tanaka, Daisuke Umeki, Shota Kido, Rikako Makishima, Yuko Tamaki, Takumi Murakami, Masayuki Fujiwara, Yusuke Saijo

**Affiliations:** Yanmar Holdings Co., Ltd., 1-32 Chayamachi Kita-ku, Osaka, Japan; Nara Institute of Science and Technology, Graduate School of Biological Sciences, 8916-5 Takayama-cho, Ikoma, Nara, Japan; School of Life Science and Technology, Tokyo Institute of Technology, 2-12-1 Ookayama, Meguro-ku, Tokyo, Japan; Advanced Genomics Center, National Institute of Genetics, 1111 Yata, Mishima, Shizuoka, Japan

**Keywords:** Biocontrol, bacterial wilt, tomato, plant-associated microbes, root microbiome, SynCom

## Abstract

Host-protective or disease-suppressive microorganisms are anticipated as sustainable controls for crop diseases, such as bacterial wilt. However, the efficacy of biocontrol strategies is often limited by a lack of resilience under varying environmental conditions and interactions with native microbial communities in the field. In this study, we assembled a bacterial wilt-suppressive synthetic community (SynCom) in tomato, consisting of bacterial isolates derived from co-cultured microbial complexes associated with the plant. This SynCom demonstrates significant disease-suppressive effects against *Ralstonia solanacearum* in tomato seedlings under both axenic and soil conditions. Additionally, our findings suggest the existence of an optimal level of SynCom colonization in plants, which is critical for effective disease control. Furthermore, the SynCom exhibits direct antibiotic activity and immunogenic properties that enhance the production of defense-related secondary metabolites, thereby influencing the plant-associated microbiome. Our results provide an effective approach to constructing SynComs that exert disease-suppressive effects within microbial community contexts.

## INTRODUCTION

It is estimated that approximately 20-40% of global food production is lost each year due to plant diseases (FAO, 2017). Among these diseases, soil-borne pathogens contribute to 10-20% of annual yield loss (Yuliar *et al*., 2015). Bacterial wilt, a devastating plant disease characterized by rapid and fatal wilting symptoms, is caused by the gram-negative β-proteobacterium *Ralstonia solanacearum* species complex (RSSC). *Ralstonia solanacearum* ranks among the top ten bacterial pathogens of economic importance in agriculture (Mansfield *et al*., 2012). The RSSC has a wide host range, infecting over 250 plant species worldwide (Prior *et al*., 2016). It primarily invades plants through the root elongation zone, sites of lateral root emergence, and physical wounds. The RSSC multiplies around the stele, filling the xylem vessels with large quantities of bacterial cells and extracellular polysaccharides, ultimately leading to significant wilting in host plants (Schell, 2000).

Biocontrol using plant-protective microbes has been proposed as a preferred method for managing RSSC infections in sustainable agricultural practices (Yuliar *et al*., 2015). Several biological control agents (BCAs) have been reported for bacterial wilt, including non-virulent strains of RSSC (Wei *et al*., 2013; Marian *et al*., 2018), *Pseudomonas* spp. (Kumatani *et al*., 2009; Singh *et al*., 2013), *Bacillus* spp. (Phae *et al*., 1992; Yuan *et al*., 2014; Fu *et al*., 2020), *Mitsuaria* spp. (Marian *et al*., 2018), *Flavobacterium* spp. (Kwak *et al*., 2018), *Trichoderma* spp. (Singh *et al*., 2012), *Acinetobacter* spp. (Xue *et al*., 2009), and *Stenotrophomonas* spp. (Messiha *et al*., 2007). These BCAs confer pathogen resistance through competition for nutrients and space, antibiotic production, and/or induction of systemic resistance in the host. Additionally, microbiome members that produce siderophores ineffective for RSSC, such as pyoverdines, can effectively suppress bacterial wilt (Gu *et al*., 2020a; 2020b). However, successful application of biological controls against soilborne diseases has been limited, partly because introduced biocontrol microbes are often outcompeted or suppressed by indigenous microbes in the field (Mihorimbere *et al*., 2011; Mazzola and Freilich, 2017).

Application of microbial communities has emerged as an alternative strategy to address these challenges. This approach allows for the combination of microorganisms with varying traits, enhancing their collective benefits and resilience in diverse habitats, including different host genotypes (Compant *et al*., 2019; Berihu *et al*., 2023). Within microbial networks, certain microbes, termed keystone species, are extensively and closely interconnected with others, playing pivotal roles in plant-microbiota interactions (Niu *et al*., 2017; Banerjee *et al*., 2018). When microbial complexes are introduced to plants during early developmental stages, they may function as keystone species or provide a foundational framework for establishing plant-associated microbiota (Trivedi *et al*., 2021; Capargo *et al*., 2023).

In this study, we investigate disease-suppressive strains of root-associated bacteria co-isolated from tomato roots that are effective against *Ralstonia solanacearum*. The synthetic communities (SynComs) formed by these strains outcompete *Ralstonia solanacearum* in tomato, providing an effective biocontrol strategy for bacterial wilt diseases. Our results further suggest the potential of this method for obtaining beneficial bacterial SynComs.

## RESULTS

### Identification of microbial complexes suppressing bacterial wilt disease

We hypothesized that disease-suppressing microbes contribute to the health of tomato plants in pesticide-free farms. To test this, we isolated microbial complexes from the roots of tomato plants grown in pesticide-free farms in Japan (Okayama, Osaka, Shiga, and Nara Prefectures). From initial bacterial cultivation, we isolated 1,934 plant-associated microbial complexes (Fig. 1A, Supplementary Table 1).

**Figure 1.**
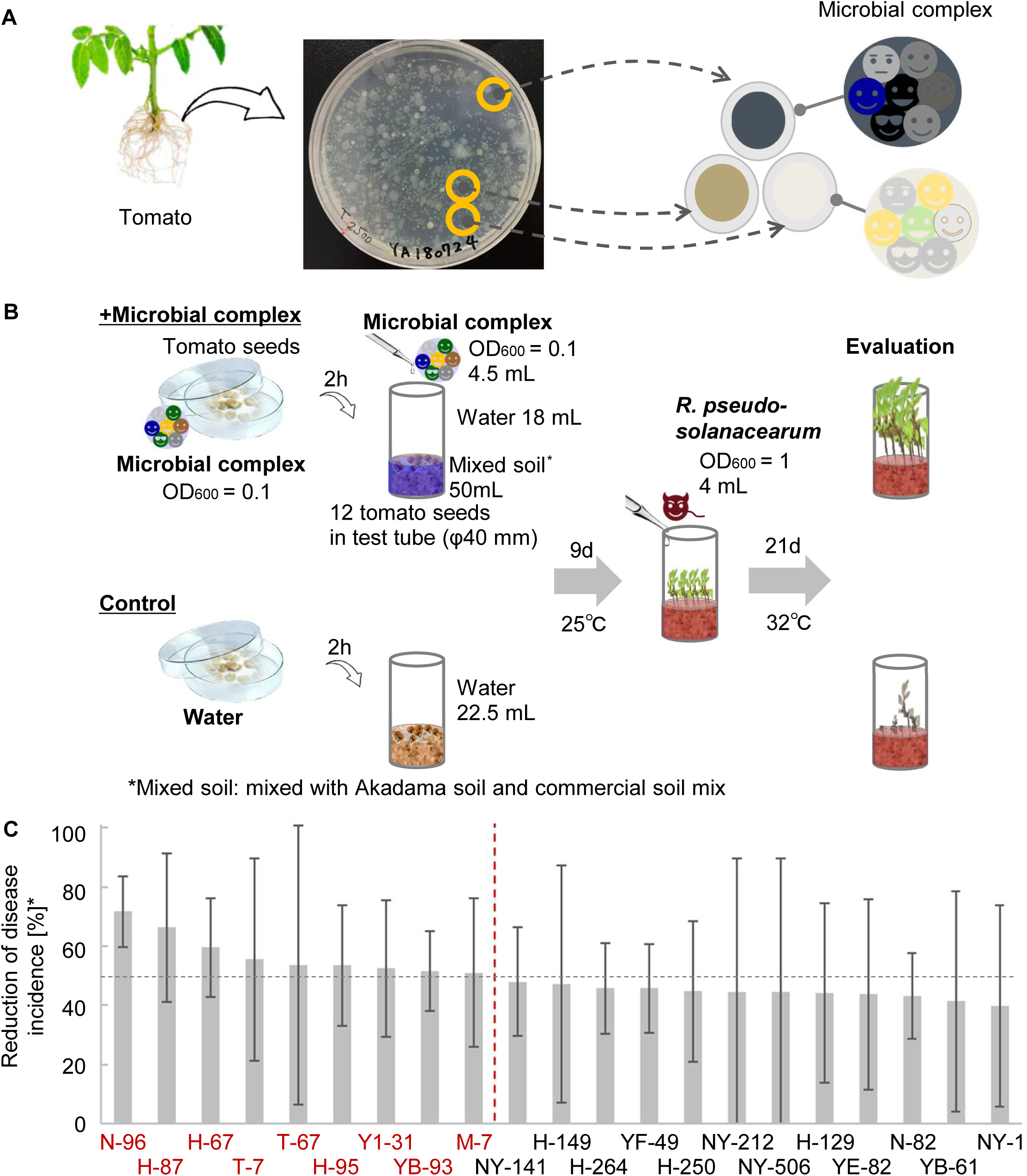
Selection of disease-suppressive microbial complexes. A) Plant-associated microbial complexes were isolated from the roots of tomato plants grown in pesticide-free farms. B) Microbial complexes were selected in a non-sterile tomato seedling bioassay. C) Nine microbial complexes reduced disease incidence by more than 50% compared to the mock controls in the tomato seedling bioassay. The graph shows that 21 microbial complexes reduced disease incidence by more than 40% compared to the mock controls. The treatments on the left side of the red vertical dashed line indicate those that reduced disease incidence by more than 50%. The black horizontal line indicates a 50% reduction in disease incidence. Error bars represent the standard deviation of the mean (the number of plants investigated was 9-12. The experiments were repeated three times).

These complexes were then screened for their ability to suppress bacterial wilt in non-sterile tomato seedlings in soil tube settings (Fig. 1B). Seeds were soaked in microbial complex suspensions for two hours before being sown in mixed soil, with the suspensions also added to the soil. Nine days later, the plants were inoculated with *Ralstonia pseudosolanacearum* strain Miho1, and the incidence of bacterial wilt was assessed 21 days post-inoculation. Nine microbial complexes reduced disease incidence by over 50% compared to the mock controls (Fig. 1C).

These nine complexes were further validated in soil pot settings (Fig. 2). Seeds soaked in the microbial complex suspension were sown in soil and inoculated again with the suspension. Twenty-one-day-old seedlings were then transplanted into fresh soil and grown for nine days before being inoculated with *R. pseudosolanacearum.* At 14 days post-inoculation, the H-87 complex showed the strongest disease-suppressive effects (Table 1), suggesting that it contains disease-suppressive bacterial isolates.

**Figure 2.**
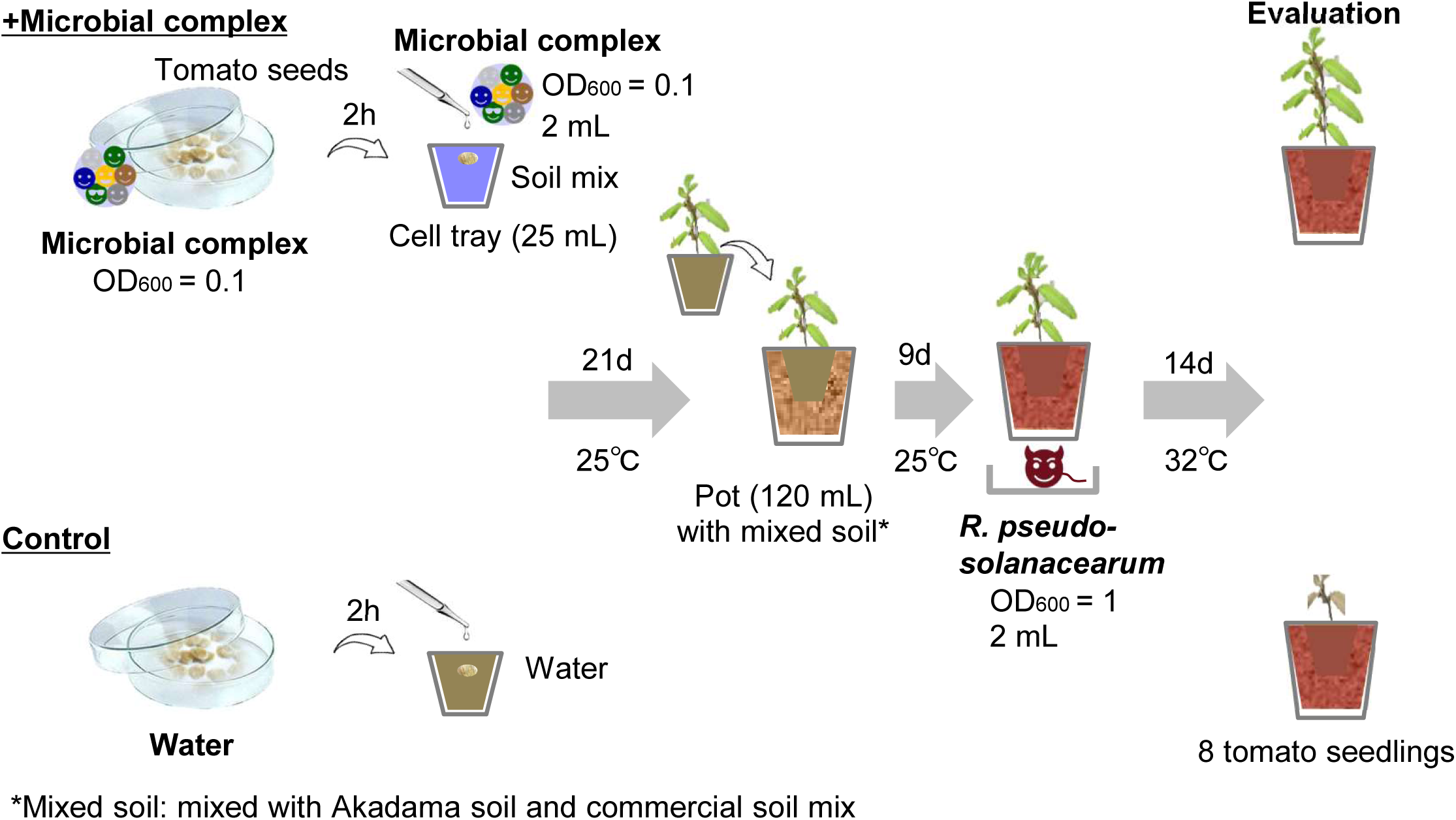
Validation of disease-suppressive effects of microbial complexes. Nine microbial complexes selected in the seedling bioassay were validated for their disease-suppressive effects on seedlings in pots.

**Table 1.** Microbial complex H-87 showed the highest disease-suppressive effects on tomato seedlings grown in pots with a potting soil mix among all the microbial complexes tested.

| Microbial<br>complex | Reduction of disease incidence [%] |  |  |  |  |
| --- | --- | --- | --- | --- | --- |
|  | 1st | 2nd | 3rd | 4th | Average |
| H-87 | 67 | 13 | -40 | 83 | 31 |
| N-96 | 33 | 33 | 0 | 16 | 21 |
| M-7 | 50 | 17 | 0 | 14 | 20 |
| Y1-31 | -81 | 14 | 100 | 16 | 12 |
| T-7 | 71 | 0 | -50 | 0 | 5 |
| YB-93 | -36 | 33 | 0 | 17 | 4 |
| H-67 | -33 | 25 | 0 | - | - |
| H-95 | -67 | 0 | 25 | - | - |
| T-67 | -8 | -50 | 0 | - | - |

The number of plants investigated was 8. The experiments were repeated three times or four times (the fourth experiment was conducted for the microbial complexes that showed disease suppression effects in three experiments). Six microbial complexes showed disease-suppressive effects on tomato seedlings grown in pots with a potting soil mix.

### Construction of bacterial wilt disease-suppressive SynCom H87-SC6

To reconstruct a SynCom with disease-suppressive activity, we isolated and examined individual bacterial strains from the H-87 microbial complex. We recovered eighty bacterial isolates from H-87 and screened them for disease-suppressive individuals in non-sterile seedlings. Seven strains (H87-6, H87-25, H87-33, H87-40, H87-50, H87-54, and H87-67) reduced disease incidence by over 10% compared to controls (Fig. 3A).

**Figure 3.**
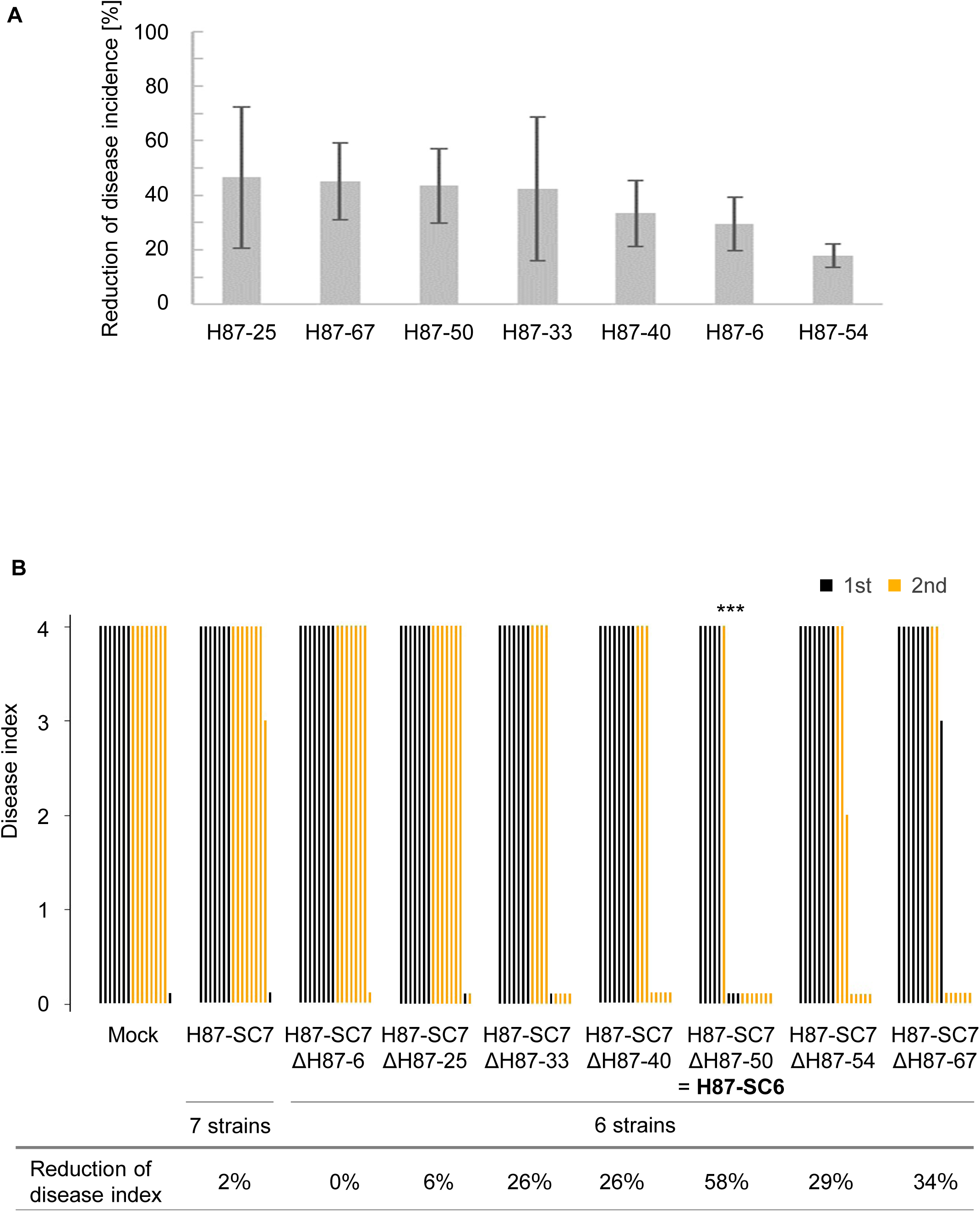
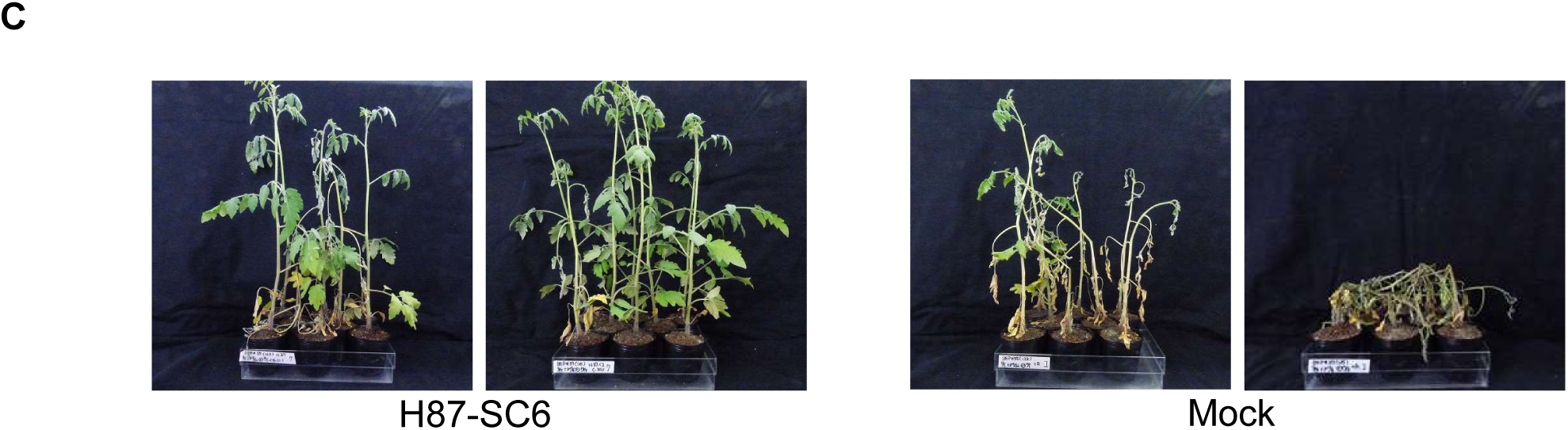
SynCom suppresses bacterial wilt disease in tomato seedlings grown in pots with a potting soil mix. A) Seven bacterial strains isolated from H-87 reduced disease incidence by more than 10% compared to the mock controls in a non-sterile tomato seedling bioassay. Error bars represent the standard deviation of the mean (the number of plants investigated was 9-12. The experiments were repeated two times). B) SynCom H87-SC6 (H87-SC7ΔH87-50) showed the greatest reduction in disease symptoms among the SynComs constructed with seven or six disease-suppressive strains in pot experiments. The disease index scale is as follows: 0; no disease symptoms, 1; up to 25% leaf wilted, 2; up to 50% leaf wilted, 3; up to 75% leaf wilted, 4; up to 100% leaf wilted. Each bar represents the disease index for an individual plant The number of plants investigated was 8 and the experiments were repeated twice. The data were subjected to one-way ANOVA with Dunnett’s multiple comparison test (***, p < 0.001 compared to the corresponding values of mock controls). C) Disease suppression phenotypes of plants treated with H87-SC6 and mock controls in pot experiments.

We then constructed a SynCom designated H87-SC7 by combining these seven strains. Additionally, we constructed 6-isolate SynComs by excluding one strain at a time from H87-SC7. In pot experiments, one of these SynComs, designated H87-SC6, which excluded H87-50, significantly reduced disease incidence (Fig. 3B, 3C), while the other 6-isolate SynComs did not show similar effects (Supplementary Fig. 1A).

Notably, the individual isolates of H87-SC6 did not suppress disease on their own in the pot experiments (Supplementary Fig. 1B), indicating that these strains exert strong disease-suppressive effects only when combined as a SynCom. This suggests that even non-suppressive isolates can confer robust disease suppression when functioning together as a community.

### Disease suppression by H87-SC6 is optimal at specific bacterial colonization levels

We investigated the possible relationship between root colonization levels and disease suppression by the SynCom H87-SC6. Surface-sterilized seeds were inoculated with H87-SC6 and grown under sterile conditions. We determined bacterial growth at 9 d post inoculation (dpi) with H87-SC6, immediately before inoculating with *R. pseudosolanacearum*, and assessed disease incidence at 12 dpi (Supplementary Fig. 2A). We found that disease incidence decreased as the *in-planta* growth of H87-SC6 increased, up to 6×10^5^ to 8×10^5^ cfu/g (plant fresh weight). However, as bacterial growth exceeded ∼10^7^ cfu/g, disease suppression diminished (Supplementary Fig. 2B). The results suggest that the disease-suppressive effects of H87-SC6 are most effective within an optimal colonization range of approximately 10^5^ to 10^6^ cfu/g.

### H87-SC6 composition and taxonomic identification

We identified the six isolates constituting H87-SC6 by sequencing their 16S rRNA genes and conducting BLASTN searches against the database Representative genomes (ref_prok_rep_genomes). The isolates were classified into three genera: H87-6 and H87-25 belong to *Leucobacter*, H87-33 to *Alcaligenes*, and H87-40, H87-54, and H87-67 to *Stenotrophomonas* (Table 2).

**Table 2.** Strains that construct H87-SC6 phylogenetically belong to the genus *Leucobacter, Alcaligenes*, or *Stenotrophomonas*. Bacterial strains were identified based on the sequences of their 16S rRNA gene. Bacterial cells were amplified by colony PCR using primers 27F (5’-AGAGTTTGATCCTGGCTCAG-3’) and 907R (5’-CCGTCAATTCMTTTRAGTTT-3’), and their identities were determined by sequencing. The obtained sequences were then subjected to homology searches using BLAST.

| Strain | Taxonomy |
| --- | --- |
| H87-6 | *Leucobacter komagatae* |
| H87-25 |  |
| H87-33 | *Alcaligenes faecalis* |
| H87-40 | *Stenotrophomonas maltophilia* |
| H87-54 |  |
| H87-67 |  |

These genera are known to include strains that benefit host plants. For example, *Leucobacter* species enhances drought tolerance in rice (Bates and King, 2021), *Alcaligenes* sp. promotes plant growth in maize under salinity stress (Fatima *et al*., 2020), and *Stenotrophomonas* sp. not only promote wheat growth (Singh and Jha, 2017) but also exhibit antibacterial activity against bacterial wilt (Elhalag *et al*., 2016).

### H87-SC5 and H87-SC3 suppress bacterial wilt disease under axenic conditions

To determine which isolates of H87-SC6 are essential for disease suppression, we tested combinations of five out of the six isolates. Surface-sterilized tomato seeds were inoculated with H87-SC6 or its 5-isolate variants and then grown under sterile conditions for 9 days in soil tubes. The plants were then inoculated with *R. pseudosolanacearum* and assessed for disease incidence at 12 dpi (Supplementary Fig. 3A).

All 5-isolate SynComs effectively suppressed disease incidence (Supplementary Fig. 3B). Notably, one combination, H87-SC5 (excluding H87-25), showed even greater disease suppression compared to H87-SC6 (Supplementary Fig. 3B). However, while these 5-isolate SynComs were effective under sterile conditions, they failed to suppress disease in non-sterile pot experiments (Supplementary Fig. 1A), suggesting that their function may be influenced by other soil-derived microbes.

Given the superior performance of H87-SC5 under sterile conditions, it appears that H87-25 may compromise the SynCom’s disease-suppressing ability in these settings. We then constructed a 3-isolate SynCom, H87-SC3, consisting of one strain each from the genera *Leucobacter*, *Alcaligenes*, and *Stenotrophomonas*. In axenic tube assays (Fig. 4A), H87-SC3 demonstrated similar disease suppression to H87-SC5 (Fig. 4B, 4C), indicating that these three isolates are sufficient to confer resistance against bacterial wilt.

**Figure 4.**
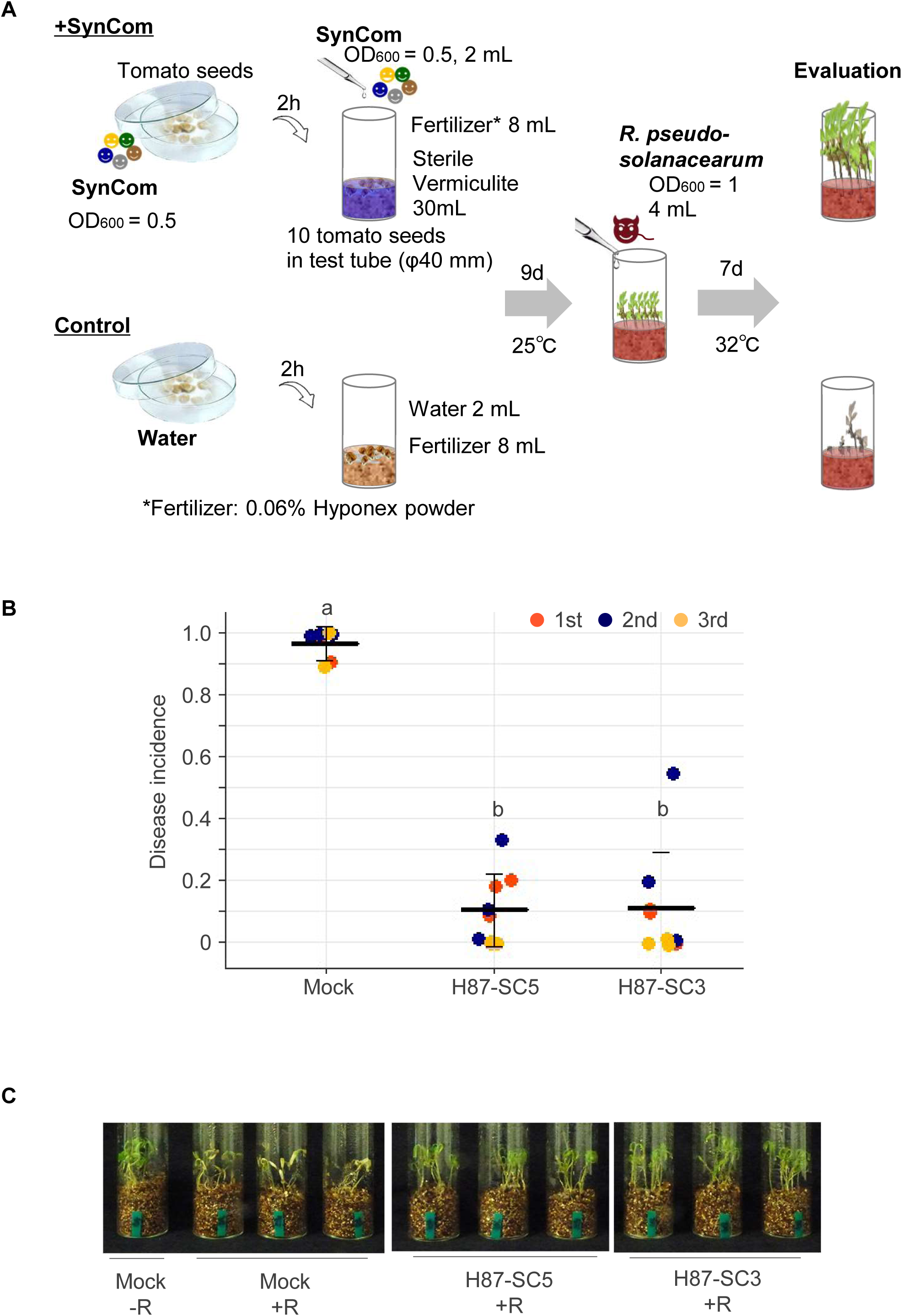
SynComs constructed with strains belonging phylogenetically to the genus *Leucobacter*, *Alcaligenes*, or *Stenotrophomonas* show disease-suppressive effects. A) Validation of the disease-suppressive effects of SynCom H87-SC5 and H87-SC3 in a seedling bioassay under sterile conditions. B) H87-SC5 (constructed with one *Leucobacter* sp., one *Alcaligenes* sp., and three *Stenotrophomonas* sp.) and H87-SC3 (constructed with one *Leucobacter* sp., one *Alcaligenes* sp., and one *Stenotrophomonas* sp.) exhibit disease-suppressive effects seven days post inoculation of *R. pseudosolanacearum*. The error bars represent the standard deviation of the mean. The middle line represents the average disease incidence (8-10 plants were investigated per tube, with three tubes per experiment. Experiments were repeated three times). The data were analyzed using one-way ANOVA with Tukey-Kramer’s multiple comparison test. Different letters indicate significant differences at p < 0.05. C) Disease suppression phenotypes of plants treated with H87-SC6, H87-SC5, H87-SC3, and mock controls seven days post inoculation of *R. pseudosolanacearum* (2nd experiment).

We determined the genome sequences of the three isolates in H87-SC3: H87-6, H87-33 and H87-40. Phylogenetic analysis revealed that H87-6 is likely a novel species of *Leucobacter*, with an average nucleotide identity (ANI) < 95% compared to existing genomes in the NCBI GenBank database (accessed on May 15th, 2023) (Fig. 5A, Supplementary Table 2). H87-33 was identified as *Alcaligenes faecalis* (ANI > 98%) (Fig. 5B), and H87-40 as *Stenotrophomonas maltophilia* (ANI > 98%) (Fig. 5C).

**Figure 5.**
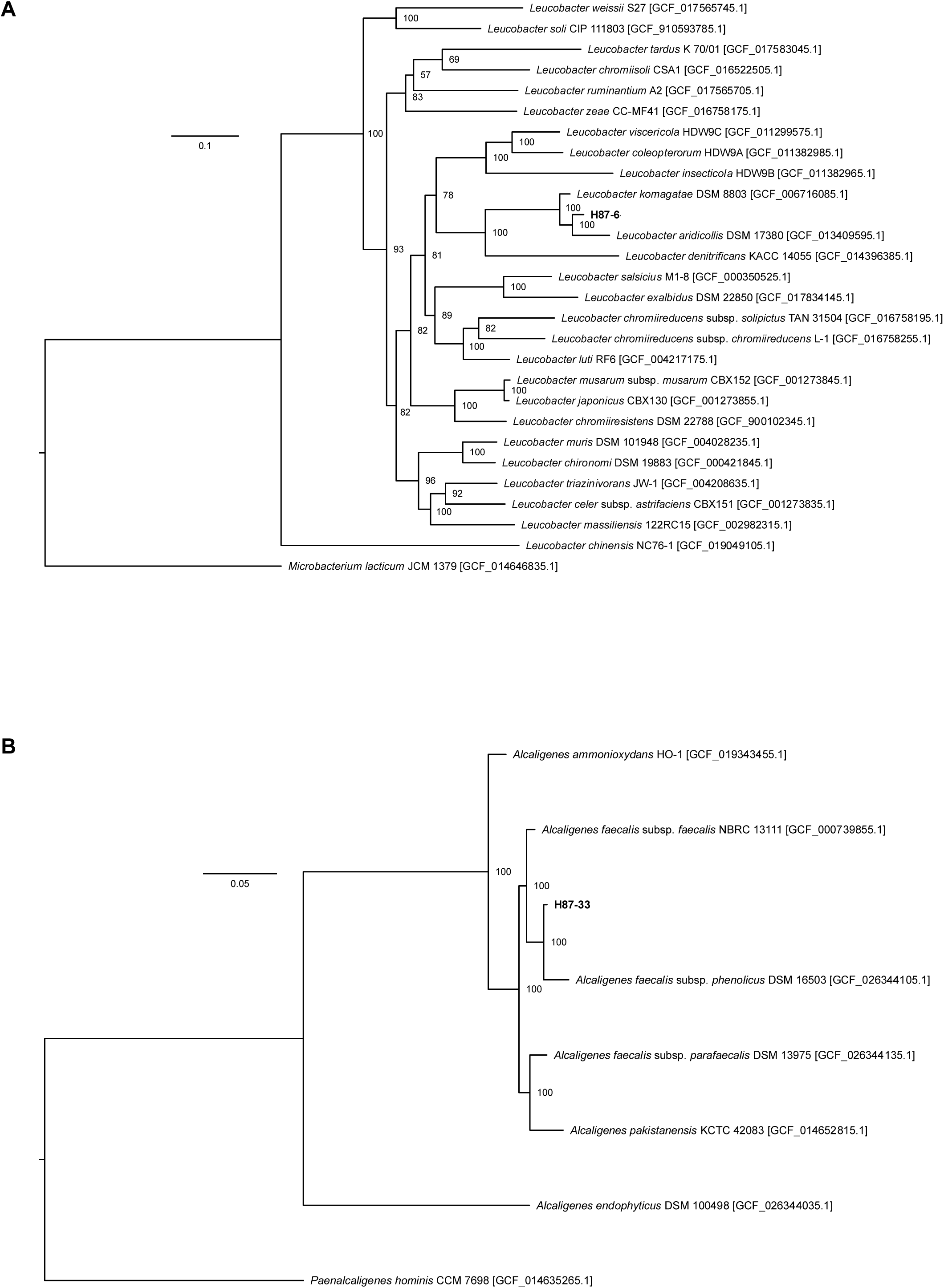
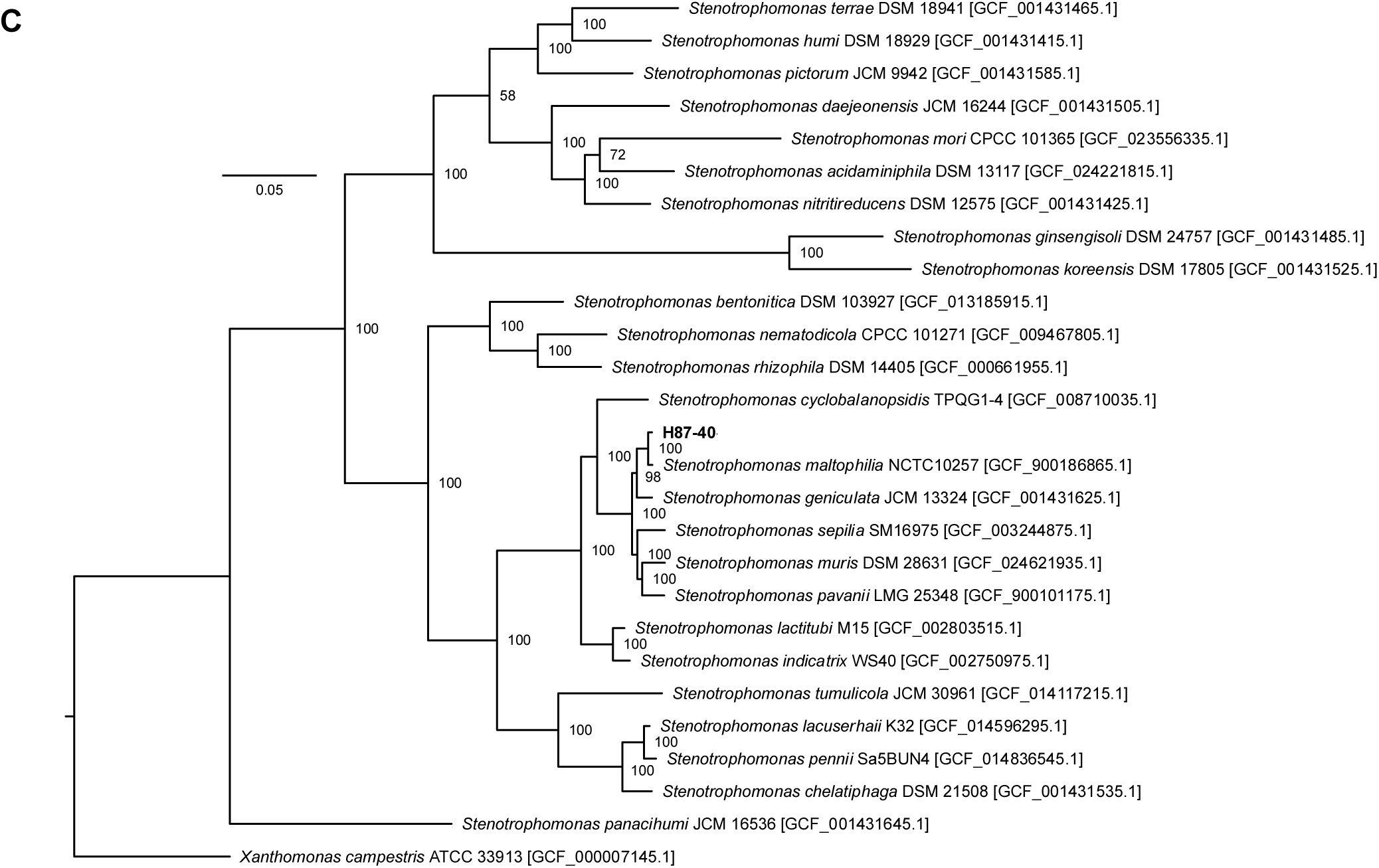
Phylogenetic positions of three strains (H87-6, H87-33, H87-40) based on the concatenated amino acid sequences of the conserved marker genes. The trees were constructed using the maximum-likelihood method with the LG+I+G4+F model and 500 bootstrap resamplings. Bootstrap confidence values >60% are displayed. The scale bar indicates the number of substitutions per site. A) The strain H87-6 is currently considered appropriate to be described as *Leucobacter* sp. H87-6. The relatively related species to H87-6 are *L. komagatae* DSM 8803^T^ (ANI: 93.9%) and *L. aridicollis* DSM 17380^T^ (ANI: 86.2%), but both fall below the ANI threshold of 95% required to be considered the same species. B) The strain H87-33 is considered to belong to at least *Alcaligenes faecalis*. H87-33 showed a high ANI value (96.2%) with *A. faecalis* subsp. *phenolicus* DSM 16503^T^. C) The strain H87-40 is considered to belong to *Stenotrophomonas maltophila*. H87-40 showed a high ANI value (98.1%) with *S. maltophila* NCTC 10257^T^.

Shoot infection by *R. pseudosolanacearum* is suppressed in plants inoculated with H87-SC5 or H87-SC3.

We investigated whether the suppression of bacterial wilt by SynComs H87-SC5 and H87-SC3 is linked to the inhibition of *R. pseudosolanacearum* growth. Tomato seeds were inoculated with H87-SC5 or H87-SC3, and 9-d-old seedlings were subsequently inoculated with *R. pseudosolanacearum.* At 5 dpi, we observed a significant reduction in *R. pseudosolanacearum* growth in the shoots of plants pre-inoculated with H87-SC5 or H87-SC3, while *R. pseudosolanacearum* growth in the roots and rhizosphere remained unaffected regardless of SynCom inoculation (Fig. 6A). Interestingly, there were no significant differences in the growth of SynCom bacteria between the shoots and the roots at 8 dpi, prior to *R. pseudosolanacearum* inoculation (Fig. 6B). These results suggest that H87-SC5 and H87-SC3 primarily suppress *R. pseudosolanacearum* infection during its root-to-shoot migration or within the shoots (Fig. 6C).

**Figure 6.**
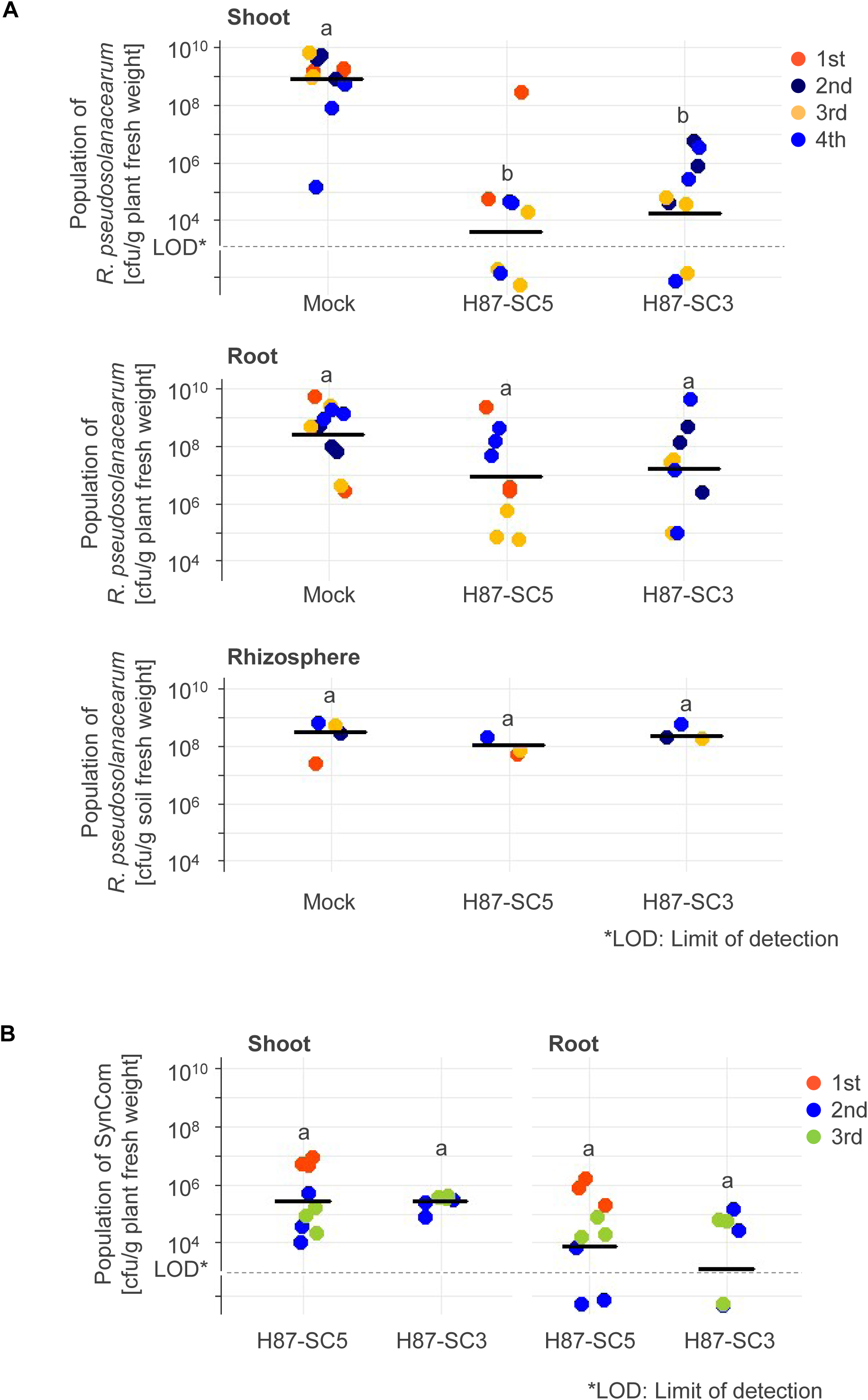
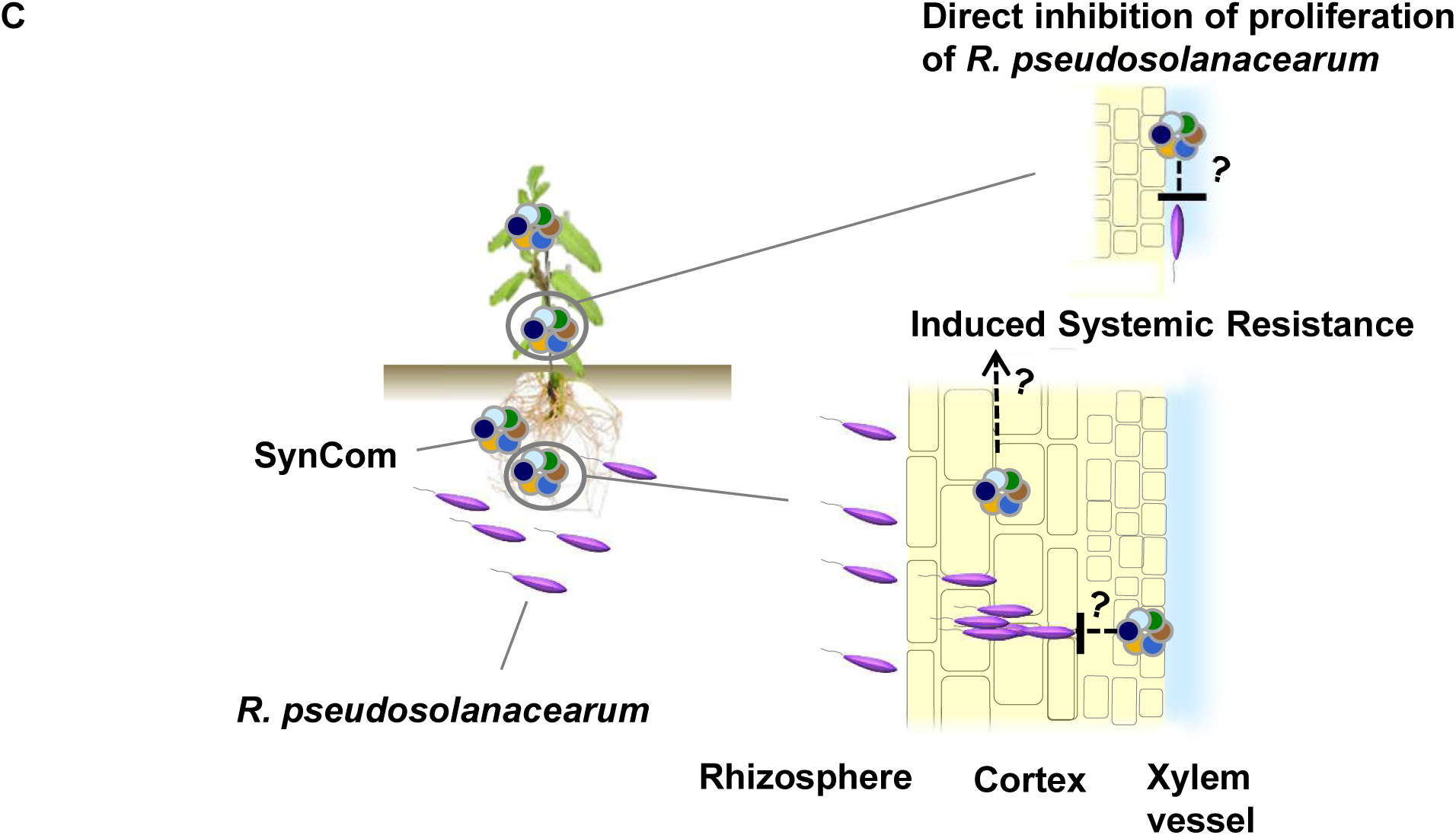
Inhibition of *R. pseudosolanacearum* proliferation, migration, or invasion by H87-SC5 or H87-SC3 in tomato plant shoots. A) The population of *R. pseudosolanacearum* in tomato plant shoots treated with H87-SC5 or H87-SC3 is lower compared to mock controls five days after inoculation with *R. pseudosolanacearum*. B) No significant differences in SynCom populations between shoot and root eight days after seeding. A) and B) The middle line represents the average population of *R. pseudosolanacearum* (n = 4 plants per experiment. The first, third, and fourth experiments were conducted for H87-SC5 treatments, while the second, third, and fourth experiments were conducted for H87-SC3 treatments). Data were analyzed using one-way ANOVA with Tukey-Kramer’s multiple comparison test. Different letters indicate significant differences at p < 0.05. C) SynCom may directly or indirectly inhibit *R. pseudosolanacearum* migration in the root, penetration into xylem vessels, colonization, or proliferation in the shoot.

### H87-SC6, H87-SC5 and H87-SC3 directly suppress *R. pseudosolanacearum*

We investigated whether the SynComs H87-SC6, H87-SC5, and H87-SC3 directly inhibit *R. pseudosolanacearum* growth in culture. After co-incubating *R. pseudosolanacearum* with the SynCom bacteria for 2 d, we assessed the antibacterial activities of SynComs by measuring the zones of *R. pseudosolanacearum* growth inhibition. All three SynComs—H87-SC6, H87-SC5, and H87-SC3—effectively suppressed *R. pseudosolanacearum* growth. Additionally, individual isolates H87-33, H87-40, H87-54 and H87-67 also showed substantial antibacterial activity (Fig. 7). In contrast, isoate T7-34, from the genus *Sphingomonas*, exhibited no antibacterial activity against *R. pseudosolanacearum*, consistent with its failure to suppress bacterial wilt in tomato plants (Supplementary Table 3). These results suggest that the disease suppression by these SynComs is partially due to direct competition between the SynCom bacterial isolates and *R. pseudosolanacearum*.

**Figure 7.**
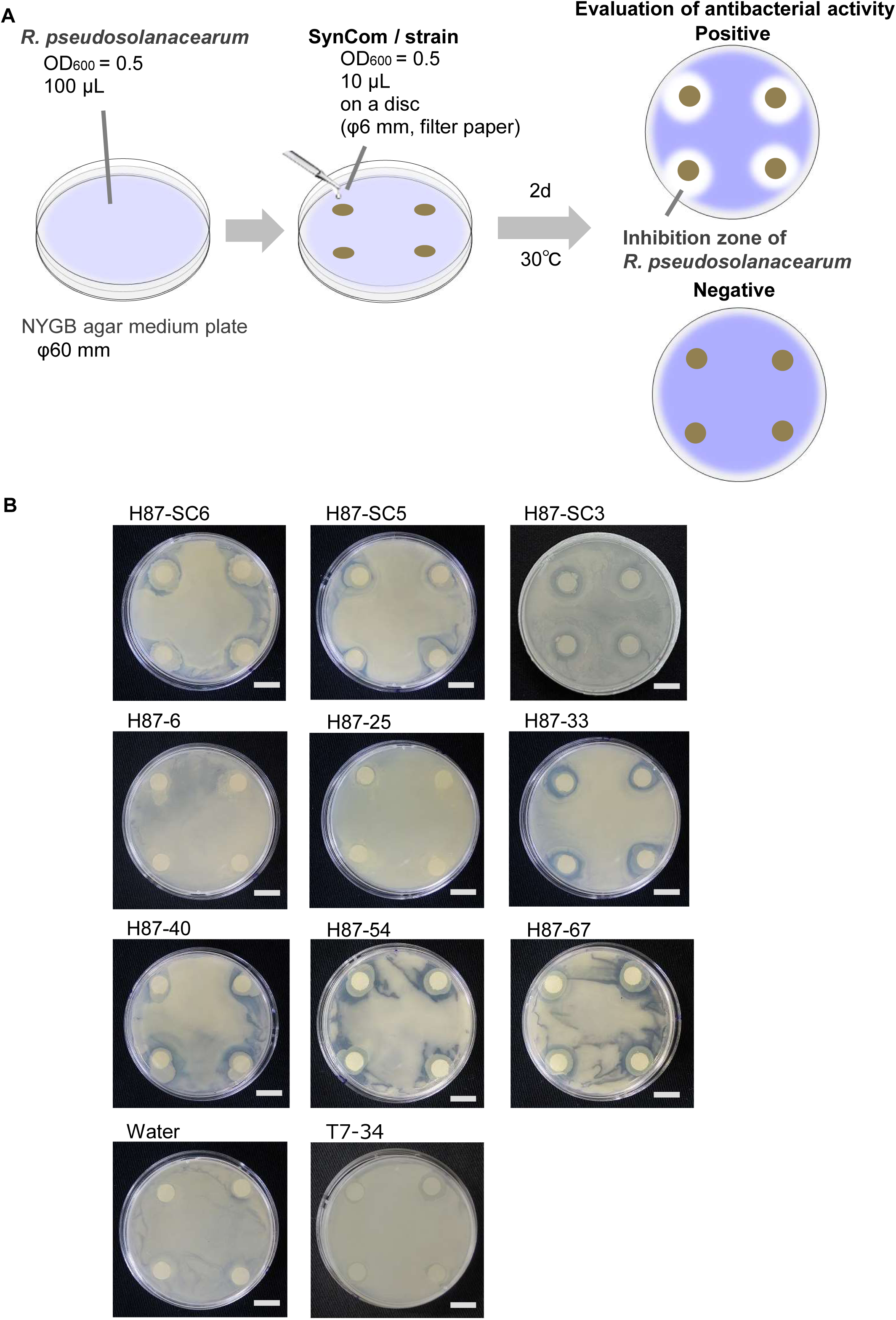
Antibacterial activity of SynComs and strains against *R. pseudosolanacearum*. A) The antibacterial activity of SynComs or strains against *R. pseudosolanacearum* was assessed using an antibacterial activity test. B) *R. pseudosolanacearum* suspension (OD_600_ = 0.5, 100 μL) was spread on the NYGB agar plate, and then SynCom or strain suspensions (OD_600_ = 0.5, 10 μL) were inoculated on filter paper discs placed on the agar plate. H87-SC6, H87-SC5, and H87-SC3 were SynComs that exhibited disease-suppressive effects against *R. pseudosolanacearum*. H87-6, H87-25, H87-33, H87-40, H87-54, and H87-64 were the strains involved in constructing the SynComs. T7-34 was the strain that did not show a disease-suppressive effect against *R. pseudosolanacearum*. The scale bar represents 10 mm.

### Possible genome basis for antibacterial functions of SynCom H87-SC3

To gain insight into the functional basis for the antibacterial activities of H87-6, H87-33, and H87-40, which constitute SynCom H87-SC3, we sequenced their whole genomes. The genomes of H87-6, H87-33, and H87-40 contained 3,101, 4,019 and 4,050 protein-coding sequences (CDS), respectively (Supplementary Table 4).

A key finding relates to the strain H87-40, closely related to *Stenotrophomonas maltophilia* (ANI > 98%), which is known to possess a type IV secretion system (T4SS) that enhances its competitive ability against other bacteria (Nas *et al*., 2019, Bayer-Santos *et al*., 2019; Supplementary Table 5). The effector proteins secreted by T4SS in *S. maltophilia* strain K279a (Nas *et al*., 2019), which have been linked to its antibacterial activity, are highly conserved in H87-40, with over 97% sequence identity in seven of nine proteins (Supplementary Table 5). Additionally, H87-40 harbors the genes for an type II secretion system (T2SS) and a putative effector protease, homologous to those of strain K279a. Proteases produced by *S. maltophilia* have been suggested to be associated with biocontrol efficacy (*Elhalag* et al., 2016). Taken together, the secretion of these effector proteins might explain H87-40’s observed antibacterial activity against *R. pseudosolanacearum* (Fig. 7).

H87-6 also possesses T4SS, while *Alcaligenes faecalis* H87-33 contains both T4SS and T2SS (Supplementary Table 5). The roles of these secretion systems in the antibacterial functions of H87-SC3 warrant future investigation.

### H87-SC5 inoculation alters plant-associated bacterial community structure

Given that H87-SC5 and H87-SC3 demonstrated disease suppression and antibacterial activities under axenic conditions, we investigated whether these effects persist in the presence of natural microbial communities. Since the first colonizers in plant-associated environments can significantly influence subsequent microbial community establishment (Trivedi *et al*., 2021), we also examined how SynCom inoculation impacts the plant-inhabiting bacterial communities.

We inoculated surface-sterilized germinated tomato seeds with H87-SC5 or H87-SC3 and grew them for 8 d in tubes filled with non-sterile soils. After inoculation with *R. pseudosolanacearum*, we assessed bacterial wilt disease incidence at 10 dpi (Fig. 8A). H87-SC5, but not H87-SC3, significantly reduced the disease index compared to the mock control (Fig. 8B). The results suggest that the inclusion of two additional isolates in H87-SC3 (forming H87-SC5) enhances robustness in disease suppression within the microbiome context.

**Figure 8.**
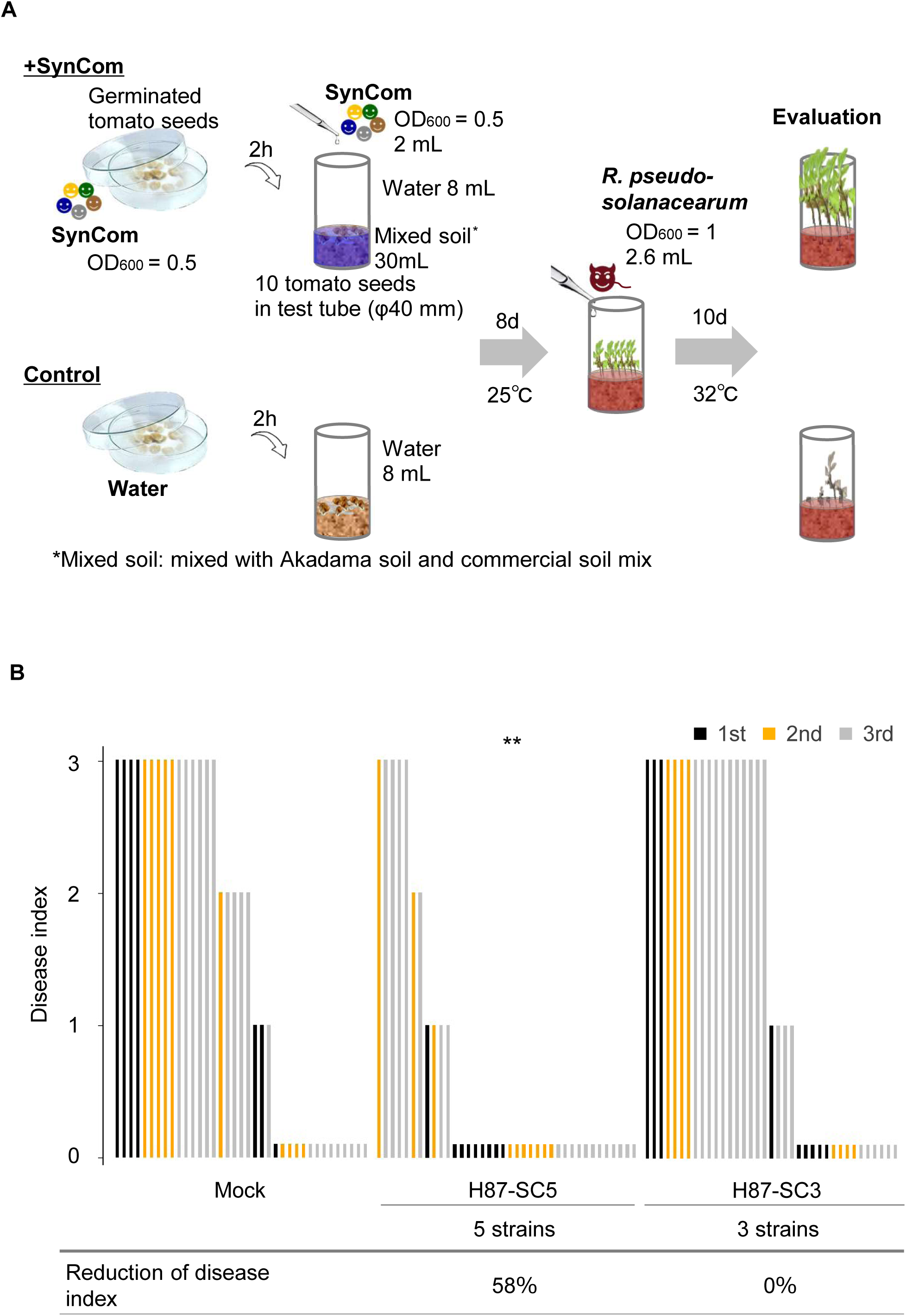
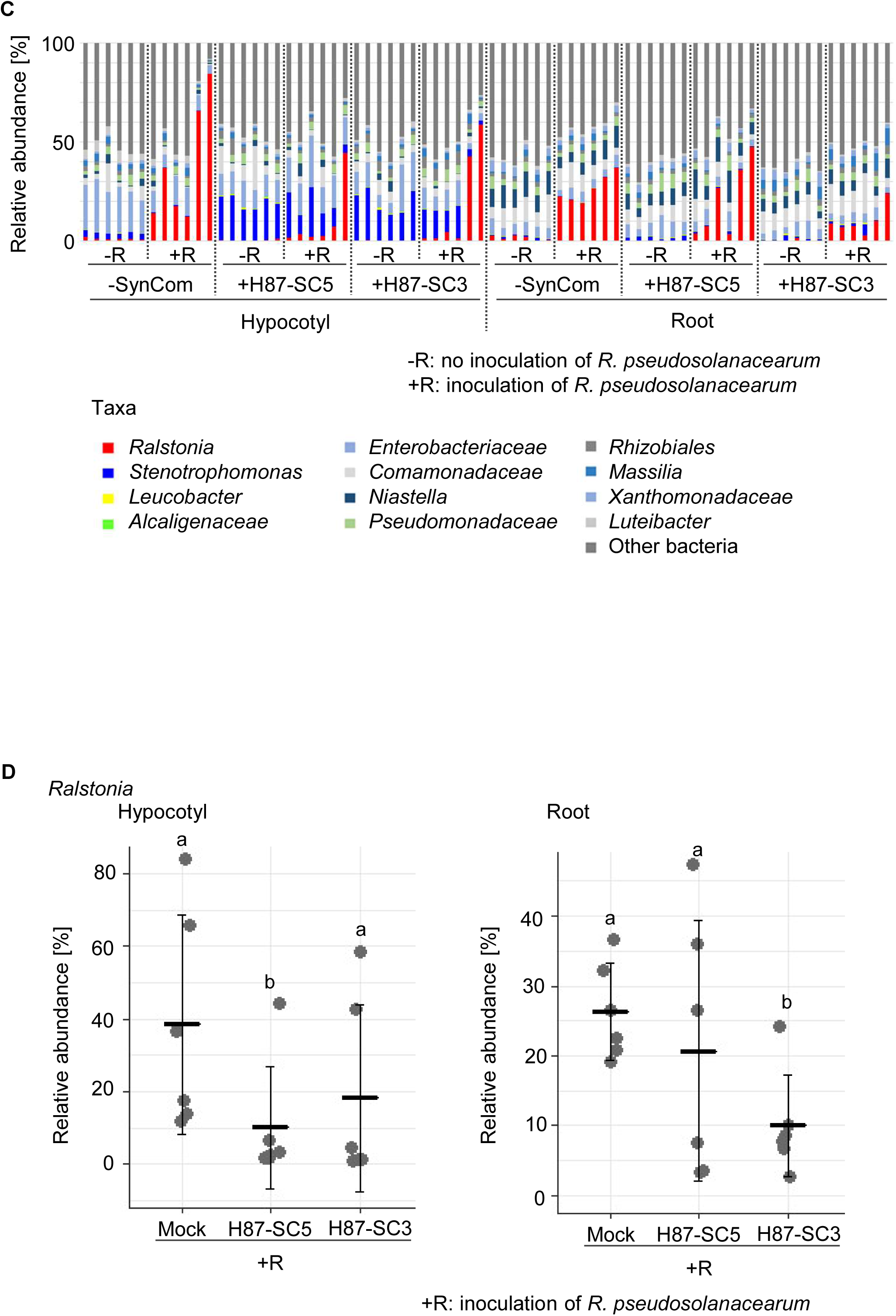
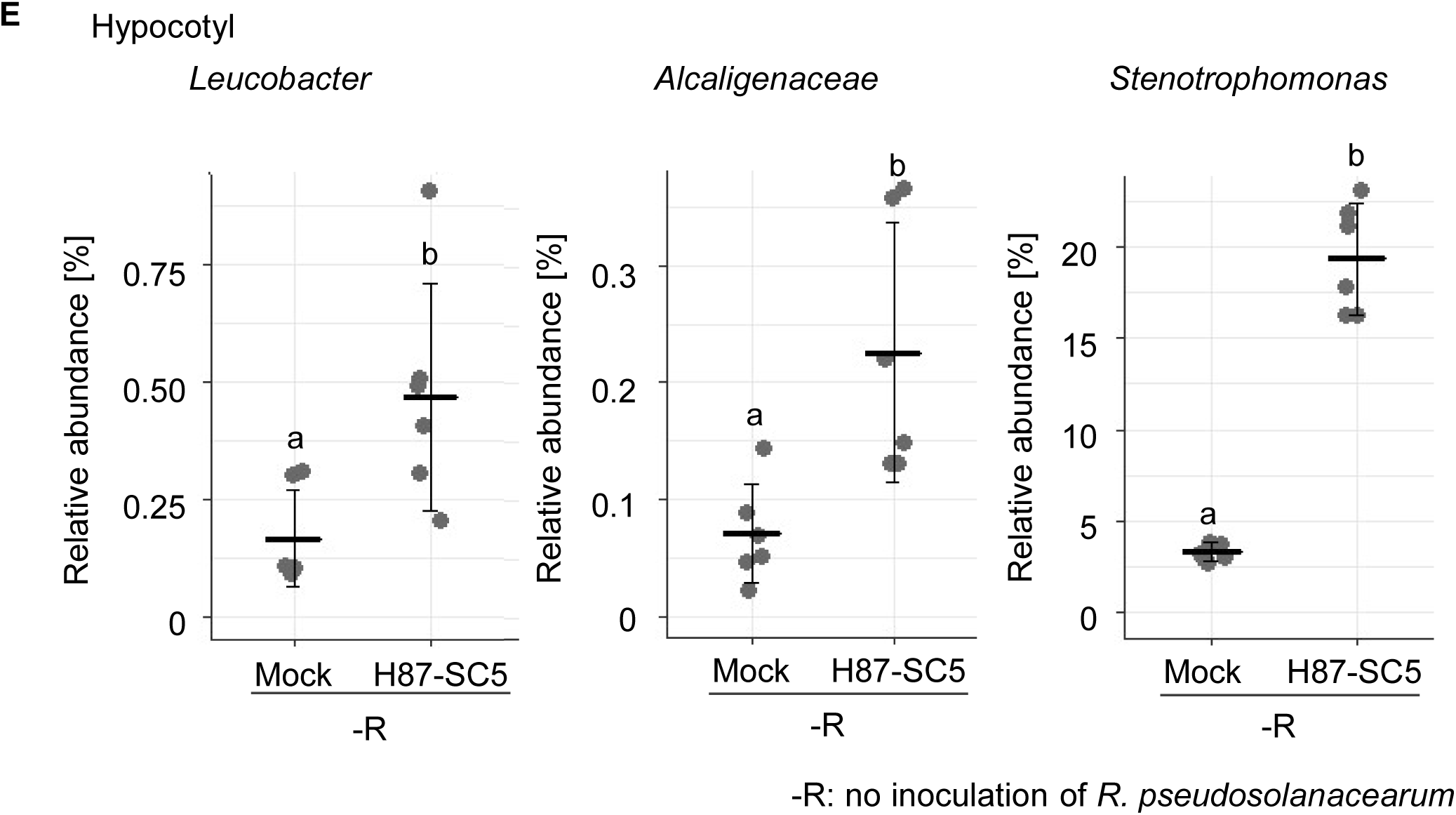
H87-SC5 shows disease-suppressive effects on tomato seedlings grown under non-sterile conditions and affects the bacterial community in plants. A) Validation of the disease-suppressive effects of SynCom H87-SC5 and H87-SC3 in a non-sterile tomato seedling bioassay. B) SynCom H87-SC5 (constructed with one *Leucobacter* sp., one *Alcaligenes* sp., and three *Stenotrophomonas* sp.) has a disease-suppressive effect 10 days post inoculation of *R. pseudosolanacearum*, while H87-SC3 (constructed with one *Leucobacter* sp., one *Alcaligenes* sp., and one *Stenotrophomonas* sp.) does not. The disease index shows: 0 - no disease symptoms, 1 - part of the leaves wilted, 2 - all leaves wilted, 3 - stem wilted. The number of plants investigated was 7-9 for the 1st test, 10 for the 2nd test, and 20 for the 3rd test per treatment. The data were subjected to one-way ANOVA with Dunnett’s multiple comparison test (**, p < 0.01 compared to the corresponding values of mock controls). C) Comparison of the bacterial community in plants grown in tubes with non- sterile mixed soil seven days post inoculation of *R. pseudosolanacearum* at the genus level. The relative abundance of taxa in each treatment and the sum of all treatments for each taon were calculated. Relative abundances were plotted for the top 37 taxa and for *Leucobacter* and *Alcaligenaceae*. For other taxa, the relative abundance was summed and designated as ‘others.’ D) Relative abundance of *Ralstonia* in plants extracted from the bacterial community data in Figure 8C. E) Relative abundance of *Leucobacter*, *Alcaligenaceae*, and *Stenotrophomonas* in plants extracted from the bacterial community data in Figure 8C. D) and E) The middle line shows the average relative abundance of the taxa (the number of samples investigated was six per treatment). Data were subjected to Kruskal-Wallis test. Different letters indicate significant differences at p < 0.05.

We then used 16S rRNA amplicon sequencing to analyze microbial community structures in the hypocotyls and roots. In mock controls, the relative abundance of *Ralstonia* in hypocotyls ranged from 12 - 84% after *R. pseudosolanacearum* inoculation (+R), whereas it was reduced to 2 - 44% in H87-SC5-inoculated plants and 1 - 59% in H87-SC3-inoculated plants (Figure 8C). H87-SC5 was more effective than H87-SC3 in limiting *Ralstonia* abundance, consistent with its stronger disease suppression (Figure 8D). Interestingly, the inoculation with H87-SC5 increased the relative abundance of beneficial bacteria such as *Leucobacter* sp., *Alcaligenaceae*, and *Stenotrophomonas* in the hypocotyls (Figure 8E). This indicates that the inoculation of H87- SC5 leads to an increase in the relative abundance of these taxa and the suppression of *Ralstonia* proliferation.

Further analysis using linear discriminant effect size (LEfSe) (Segata *et al*., 2011) revealed that H87-SC5 significantly altered the relative abundance of several bacterial taxa in both hypocotyls and roots. Specifically, H87-SC5 inoculation (-R) increased the relative abundance of 27 taxa in the hypocotyls and 17 taxa in the roots, while decreasing that of 21 and 7 taxa, respectively (Supplementary Fig. 4A). Notably, following *R. pseudosolanacearum* inoculation, *Enterobacteriaceae*, *Sphingobacterium* and *Flavobacterium* increased in the relative abundance in the hypocotyls of H87-SC5-inoculated plants (+R). Notably, *Leucobacter* and *Alcaligenaceae* decreased in abundance, but *Stenotrophomonas* remained unaffected in the hypocotyls of H87- SC5-treated plants (+R) (Supplementary Fig. 4B). The relative abundance of these taxa (*Leucobacter*, *Alcaligenaceae*, and *Stenotrophomonas*) was not significantly different in the hypocotyls between H87-SC5 and H87-SC3 inoculation. The increase in the number of the taxa was greater after H87-SC5 inoculation compared to H87-SC3 inoculation (Supplementary Fig. 4C). Principal coordinate analyses (PCoA) (in the absence of *R. pseudosolanacearum*) indicate that SynCom inoculation notably impacted the structure of plant-associated microbial communities, particularly in the hypocotyls (Supplementary Fig. 5A). Shannon’s index showed no significant differences in microbial community diversity among the mock control, H87-SC5 and H87-SC3 inoculations (Supplementary Fig. 5B), suggesting that SynCom treatment does not affect alpha diversity, at least in the absence of *R. pseudosolanacearum*. By contrast, following *R. pseudosolanacearum* inoculation, *Enterobacteriaceae*, *Sphingobacterium* and *Stenotrophomonas* decreased, while *Flavobacterium* remained unaffected, in relative abundance in the hypocotyls without SynCom inoculation (+R) (Supplementary Fig. 4D).

Overall, the results suggest that H87-SC5 not only suppresses bacterial wilt but also modulates the plant-associated microbial community, potentially contributing to disease control. Further studies are required to elucidate the specific roles of these community shifts in disease suppression.

### H87-SC6 and H87-SC5 do not inhibit tomato plant growth or yield

We assessed the impact of SynComs H87-SC6 and H87-SC5 on tomato plant growth and yield. Seedlings inoculated with these SynComs were grown in vinyl pots for 24 days, after which they were transplanted into a field free of *R. pseudosolanacearum* (Supplementary Fig. 6A).

Neither H87-SC6 nor H87-SC5 affected plant height in either pot or field settings (Supplementary Fig. 6B-C). This indicates that these SynComs do not negatively influence tomato plant growth or yield, consistent with their minimal impact on the commensal microbial communities described earlier.

## DISCUSSION

In this study, we demonstrated that applying microbial communities, specifically SynCom H87- SC6, offers an effective approach to confer bacterial wilt resistance in tomato. Additive functional benefits can be expected when combining microbes of different traits (Compant *et al*. 2019). Unlike the conventional method of artificially combining different microbial isolates, our approach utilized SynCom H87-SC6, which was co-isolated directly from tomato roots. This approach may reduce potential antagonism and competition among bacterial strains that can arise when optimal combinations are not achieved, a limitation often seen in artificially constructed SynComs (De Vrieze *et al*., 2018).

Interestingly, the addition of H87-50 to H87-SC6 to create H87-SC7 did not enhance disease suppression, suggesting that H87-50 negatively affects the disease-suppressive function of H87- SC6. It would be valuable to investigate whether H87-50 inhibits the growth or functionality of other isolates within the SynCom. Conversely, H87-SC5 (excluding H87-25) had no effect in non-sterile pot experiments but was effective in in non-sterile tube assays. Further research is needed to elucidate how H87-25 causes these differences, which may stem from the colonization states of SynCom or variations in plant-SynCom interactions due to different plant growth stages.

H87-SC6 strains of *Leucobacter*, *Alcaligenes*, and *Stenotrophomonas* were not detected in the root microbial communities of plants grown in the soil, from which they were isolated, as indicated by 16S rRNA sequencing analyses (Supplementary Table 6). This suggests that these isolates are initially low in abundance but that co-cultivation with partner bacteria effectively promotes their growth. Our co-cultivation and co-isolation methods may enhance SynCom functionality and increase isolation chances.

The population dynamics of beneficial bacteria during bacterial wilt suppression have been documented (Kwak *et al*., 2018, Fu *et al*., 2020), but the relationship between disease-suppressive and pathogenic bacteria remains underexplored. Our findings indicate that an optimal SynCom colonization range is critical for effective disease control, highlighting the importance of regulating SynCom levels.

*R. pseudosolanacearum* invades the plant through the root elongation zone, lateral root emergence sites, and root wounds, colonizing the root cortex and eventually migrating to the xylem, leading to plant collapse due to extracellular polysaccharide production that prevents xylem water transport (Hikichi *et al*., 2017). In this study, we show that SynCom inoculation reduces *R. pseudosolanacearum* populations in the shoots, although its root and rhizosphere populations were unaffected. The precise stage at which SynCom suppresses bacterial wilt—whether in migration, colonization, or proliferation - remains unclear, possibly involving antibiotic activity against the pathogen. Additionally, SynCom alters plant-associated microbial communities, notably increasing *Flavobacterium*, which has been linked to bacterial wilt suppression (Kwak, 2018). A *S. maltophilia* strain, closely related to H87-40, is known to degrade 3-hydroxy palmitic acid methyl ester (3OH-PAME), a quorum-sensing molecule in *R. pseudosolanacearum*, and can induce systemic acquired resistance (Achari & Ramesh, 2015; Elhalag et al., 2016). It is conceivable that SynCom suppresses bacterial wilt through both direct and indirect mechanisms, warranting further investigation into its effects on plant immune responses.

The SynComs developed in this study demonstrated suppressive effects on bacterial wilt disease under standardized laboratory conditions. The SynCom introduction to plants at an early growth stage results in alterations in root and hypocotyl microbial communities, likely promoting the establishment of a beneficial microbial community, as suggested by ’priority effects’ (Trivedi *et al*., 2021). To further validate these findings, it will be crucial to assess the disease control efficacy of SynComs under varying biological and physicochemical soil conditions. Successful implementation of these SynComs could play a significant role in advancing sustainable agriculture.

## MATERIALS AND METHODS

### Isolation of tomato plant-associated microbial complexes

Plant-associated microbial complexes were isolated from the roots of tomato plants grown in pesticide-free farms (Figure 1A). The tomato cultivars used were Frutica, Chika, Momotaro-8, Green Save, Kyoryoku-beiju, and Ganbarune-forte (Supplementary Table 1). Soil adhering to the roots was removed with running tap water, and then the roots were surface sterilized in 70% (v/v) ethanol for 2 minute, followed by 1% (v/v) sodium hypochlorite for 1 minutes, and rinsed with sterile distilled water. The root samples were individually homogenized in sterile distilled water (SDW) using a sterile mortar and pestle, and 6-fold serial dilutions were then prepared. 100 μL of the homogenate dilutions were spread onto the surface of agar medium (Supplementary Table 78). Colonies that appeared after 3 days to 4 weeks of incubation at 25 °C were individually stored in the presence of 15% glycerol at -80 °C until used as plant-associated microbial complexes.

Bacterial cells in Microbial complex H-87 were spread onto NYGB agar media and incubated at 25 °C for 2 days. Bacterial single colonies on the plates were transferred to fresh NYGB agar media, purified through repetitions of streaking and colony transfer. Liquid culture from purified single colonies was stored in the presence of 15% glycerol at -80 °C until used.

### Plant materials

A susceptible tomato cultivar to bacterial wilt, Chika (Takii Seed Co., Japan), was used throughout this study. Tomato seeds were surface sterilized in 70% ethanol for 1 minute, followed by 1% sodium hypochlorite for 30 minutes, and then rinsed three times with SDW.

### Non-sterile tomato seedling bioassay for select microbial complexes and bacteria

Plant-associated microbial complexes and bacteria that exhibit disease-suppressive effects against bacterial wilt were selected using a non-sterile tomato seedling bioassay. First, a microbial complex was pre-cultured on a medium plate, which was the same type of medium used for its isolation, at 25 °C for 3 days. The cells of the complex were then transferred to a medium broth of the same type and cultured at 28 °C with shaking at 180 rpm for 2 days. The culture broth was centrifuged at 2,000 rpm for 15 minutes, and the cells were washed twice with sterile distilled water (SDW). The washed microbial complex cells were suspended in SDW, and the optical density at 600 nm (OD600) was adjusted to 0.1. Surface-sterilized tomato seeds were soaked in the suspension. Twelve seeds were sown in a tube containing 50 mL of mixed soil, which consisted of 30 g of Akadama soil and 5 g of soil mix (H-150, Yanmar Co., Japan). Then, 4.5 mL of the microbial complex suspension and 18 mL of SDW were added to the mixed soil, and the plants were grown at 25 °C for 9 days prior to pathogen inoculation. The control plants were treated with SDW instead of the microbial complex suspension. The avirulent strain of *R. pseudosolanacearum* Miho1 (provided by NARO Genebank, Japan) used in this study was grown at 30 °C for 24 h on a 2,3,5- triphenyltetrazolium chloride (TTC) medium (1% peptone, 0.5% glucose, 0.1% casein hydrolysate, 1.8% agar, and 0.05% 2,3,5-triphenyltetrazolium chloride) (Kelman, 1954). The bacterial cells were then suspended in casamino acid-peptone-glucose (CPG) broth (1% peptone, 0.5% glucose, 0.1% casein acids) (Kelman, 1954) and cultured at 30°C with shaking at 200 rpm for 24 h. The bacterial cell density was diluted to an OD600 of 1 (approximately 8 × 10^8^ cfu/mL). Nine days after seeding, 4 mL of *R. pseudosolanacearum* suspension was inoculated into tubes containing the tomato plants grown in mixed soil. The plants were then grown in a growth chamber at 32 °C, 80% relative humidity, with a 12-hour light and 12-hour dark photoperiod. The incidence of bacterial wilt was examined 21 days after the inoculation of *R. pseudosolanacearum* (Figure 1B). The disease incidence and the reduction of disease incidence were calculated using the following formula: Disease incidence = (number of plants with wilt symptoms)/(number of plants investigated). Reduction of disease incidence = ((mean of disease incidence in mock controls) - (mean of disease incidence in microbial complex treatments))/(mean of disease incidence in mock controls) × 100.

### Examination of wilt resistance in pot experiments

To validate the disease-suppressive effects against bacterial wilt, the microbial complexes selected in the non-sterile tomato seedling bioassay and the bacterial strains that constructed SynComs were subjected to a pot experiment. Surface-sterilized tomato seeds were soaked in the microbial suspension as described above. The seeds were then sown in plastic cell trays (30 × 30 mm, 25 mL) containing commercial soil mix, with one seed per tray. Afterward, 2.5 mL of vermiculite and 2 mL of microbial suspension were added to the soil mix. As a control, SDW was used instead of the microbial suspension. The tomato seedlings, at the three- to four-leaf stage, were grown on plastic cell trays at 25 °C for 21 days and then transplanted into vinyl pots (6 cm in diameter) containing 90 mL of mixed soil. Nine days post-transplantation, a 2.1 mL suspension of *R. pseudosolanacearum* (prepared as described above, OD_600_ of 1) was inoculated into the tomato plants through soil drenching inoculation. The plants were then grown in a growth chamber at 32 °C and 80% RH. Fourteen days post-inoculation of *R. pseudosolanacearum*, the symptoms of bacterial wilt were examined based on disease incidence or disease index (Figure 2), using the following scale: 0, no wilting; 1, 1-25% of leaves wilted; 2, 26-50% of leaves wilted; 3, 51-75% of leaves wilted; 4, 76-100% of leaves wilted (Roberts *et al*., 1988). The reduction of disease index was calculated using the following formula: Reduction of disease index [%] = ((mean of disease index in mock controls) - (mean of disease index in SynCom treatments))/(mean of disease index in mock controls) × 100

### Construction of SynComs by combining selected bacterial strains

SynComs were constructed by combining selected bacterial strains, with each strain’s optical density individually adjusted. Bacterial strains were pre-cultured on NYGB agar plates at 25 °C for 4 days. They were then transferred to NYGB broth and cultured at 28 °C with shaking at 200 rpm for 1 day. The culture broth was centrifuged at 2,000 rpm for 15 minutes, and the cells were washed twice with SDW. The washed bacterial cells were suspended in SDW, and their optical density at 600 nm (OD600) was individually adjusted to 0.1 or 0.5. The adjusted bacterial suspensions were mixed to construct the SynComs.

### Tomato seedling bioassay under sterile conditions

The disease-suppressive effects of SynComs under sterile conditions were validated using tomato seedlings grown on sterile vermiculite. Surface-sterilized tomato seeds were soaked in the SynCom suspension (OD600 of 0.5, prepared as described above). Ten seeds were sown in a tube with 28 mL of sterile vermiculite (5 g) containing 8 mL of liquid fertilizer (0.06% Hyponex fertilizer, Fine Powder, Hyponex Japan Co., Japan), and covered with 2 mL of sterile vermiculite. Then, 2 mL of the SynCom suspension was added to the vermiculite, and the plants were grown at 25 °C before pathogen inoculation. The control plants were treated with sterile distilled water (SDW) instead of the SynCom suspension. Nine days after seeding, 2.6 mL of *R. pseudosolanacearum* suspension (OD600 of 0.4-1, prepared as described above) was inoculated into the tubes in which the tomato plants were grown by adding it to the vermiculite. Then, the plants were grown in a growth chamber at 32 °C with a 12-hour light and 12-hour dark photoperiod. The incidence of bacterial wilt was examined 10-12 days after inoculation of *R. pseudosolanacearum* (Figure 4A).

### Identification of bacterial taxonomy

Selected bacterial strains were identified based on the sequences of their 16S rRNA gene. Bacterial cells were amplified by colony PCR with primers 27F (5’- AGAGTTTGATCCTGGCTCAG-3’) and 907R (5’-CCGTCAATTCMTTTRAGTTT-3’). The amplified products were purified by agarose gel electrophoresis. The purified products were further identified by sequence determination using the primers described above, BigDye Terminator (Applied Biosystems, USA), and 3130 Genetic Analyzer (Applied Biosystems, USA). Homology searches were performed for the obtained sequences to assess similarities using the Basic Local Alignment Search Tool (BLAST) of the US National Center for Biotechnology Information (NCBI) (BLASTN search) against the database Representative genomes (ref_prok_rep_genomes).

### Whole-genome sequencing

The bacterial strains that constructed H87-SC3 were subjected to whole-genome sequencing. Genomic DNA was extracted from the bacterial strains using the NucleoSpin@ Microbial DNA kit (Macherey-Nagel, Germany). Whole-genome sequence analyses were performed at Bioengineering Lab. Co., Japan. The extracted genomic DNA was purified using AMPure XP (BECKMAN COULTER) and sheared using a g-tube (Covaris). Libraries for PacBio sequencing were generated using the SMRTbell Express Template Prep Kit 2.0 (PacBio) and subjected to long-read sequencing on the Sequel IIe (PacBio) using the Binding kit 2.2 (PacBio). Using SMRT Link (v10.1.0.119528) (PacBio), adapter sequences were removed from the sequenced reads, and subreads were generated. After aligning these subreads, consensus sequences (CCS) were created, and CCS reads with an average quality value of less than 20 per read were removed, resulting in the generation of HiFi reads. Subsequently, with Filtlong ver 0.2.0 (https://github.com/rrwick/Filtlong), HiFi reads with a length of 1000 bases or less were removed. The high-quality HiFi reads were then assembled using the default conditions of Flye ver 2.9 (Kolmogorov *et al*., 2019). The integrity of the assembled genome data was confirmed using CheckM ver 1.2.0 (Parks *et al*., 2015). Finally, genes were predicted from the assembled sequences using Prokka ver 1.14.5 (Seemann, 2014).

Average nucleotide identities (ANI) between the obtained genome sequences and those registered in the NCBI GenBank database (accessed on May 15th, 2023) were calculated using FastANI v1.33 (Jain *et al*., 2018). Phylogenetic trees were constructed using concatenated amino acid sequences of conserved marker genes of the three bacterial strains and the type strains of the genus *Leucobacter*, *Alcaligenes*, and *Stenotrophomonas*, whose genomes were registered in the NCBI RefSeq database. PhyloPhlAn v3.0.67 (Asnicar *et al*., 2020). was used for marker gene selection, alignment, and concatenation. RAxML-NG v1.1.0 (Kozlov *et al*., 2019) and the LG+I+G4+F model was used to generate the maximum-likelihood trees from concatenated sequences. The reliability of the phylogenetic tree was estimated with 500 bootstrap runs.

### Measurement of plant-associated bacterial population by colony counting

Plants that were grown under sterile conditions for eight days after sowing were used to measure the population of plant-associated bacteria. Plant samples were surface sterilized in 70% (v/v) ethanol for 30 seconds. The samples were then homogenized and serially diluted. Dilutions of homogenates were spread onto the surface of NYGB plates (Supplementary Table 7). The number of plant-associated bacterial colonies was counted after cultivation at 25 °C for 48 hours, and the population of bacterial strains was calculated as cfu/g of plant fresh weight.

### Measurement of *R. pseudosolanacearum* population by colony counting

Plants that were grown for nine days after sowing were used to measure the population of *R. pseudosolanacearum*. Plant samples were surface sterilized in 70% (v/v) ethanol for 30 seconds. The samples were homogenized and then serially diluted. Dilutions of homogenates were spread onto the surface of modified semi-selective medium, South Africa (SMSA) plates (French *et al*., 1995). The number of *R. pseudosolanacearum* colonies was counted after cultivation at 30°C for 48 hours, and the population of bacterial strains was calculated as cfu/g plant fresh weight.

### Antibacterial activity test against *R. pseudosolanacearum*

A suspension of *R. pseudosolanacearum* (OD_600_ = 0.5, 100 μL) was spread on NYGB agar medium plates (φ60 mm). Four discs (φ6 mm, filter paper) were then placed on the NYGB agar medium plates. SynCom or strain suspensions (OD_600_ = 0.5, 10 μL) were inoculated onto the discs. The plates were incubated for two days at 30°C. Antibacterial activity was examined by measuring the growth inhibition zone of *R. pseudosolanacearum*.

### Verification of no inhibition of plant growth by SynComs

SynCom H87-SC6 and H87-SC5 were subjected to a field experiment to validate that there was no inhibition of plant growth. The tomato cultivar TY Chika (Takii Seed Co., Japan) was used in this experiment. Surface-sterilized tomato seeds were soaked in the SynCom suspension (OD_600_ of 0.5, prepared as described above). The seeds were sown in plastic cell trays (30 × 30 mm, 25 mL) with commercial soil mix (one seed per tray) and covered with 2.5 mL of vermiculite. Then, 2 mL of microbial suspension was added to the soil mix. The control plants were treated with sterile distilled water (SDW) instead of the microbial suspension. Tomato seedlings (in the three- to four-leaf stage) grown in plastic cell trays at 25°C for 21 days were transplanted into vinyl pots (9 cm in diameter, 300 mL) containing soil mix. Twenty-four days post-transplantation into pots, the seedlings were transplanted into a field (Supplementary Fig. 6A). The inhibition of plant growth by SynComs was examined 32 days post-transplantation into the field, based on the plant heights.

### Non-sterile tomato seedling bioassay to validate disease-suppressive effects of SynComs

The disease-suppressive effects of the SynComs obtained were validated against bacterial wilt using a non-sterile tomato seedling bioassay. Pre-germinated tomato seeds were soaked in the SynCom suspension (prepared as described above) with an OD_600_ of 0.5. Ten seeds were then sown in a tube containing 30 mL of mixed soil (18 g of Akadama soil mixed with 3 g of soil mix (H-150, Yanmar Co., Japan)). To the mixed soil, 2 mL of the SynCom suspension and 8 mL of sterile distilled water (SDW) were added. The plants were grown at 25 °C until pathogen inoculation. As a control, the plants were treated with SDW instead of the microbial complex suspension. Eight days after seeding, a 2.6 mL suspension of *R. pseudosolanacearum* (prepared as described above) with an OD_600_ of 1 was inoculated into the tubes where the tomato plants were grown. The suspension was added to the mixed soil. The plants were then grown in a growth chamber at 32 °C, 80% relative humidity, with a 12-hour light and 12-hour dark photoperiod. The incidence of bacterial wilt was examined 10 days after inoculation with *R. pseudosolanacearum* (Figure 8A).

### 16S rRNA amplicon sequencing

For the analysis of the bacterial community, tomato seedlings treated with SynComs and grown under non-sterile conditions were subjected to 16S rRNA amplicon sequencing. The plants were surface sterilized as described above. One sample of hypocotyls or roots was collected from each of the four plants. Genomic DNA was extracted from the bacterial strains using the NucleoSpin@ Soil kit (Macherey-Nagel, Germany). A portion of the 16S rRNA genes was amplified using the primer pair (515F-TCG TCG GCA GCG TCA GAT GTG TAT AAG AGA CAG GTG CCA GCM GCC GCG GTAA and 806R-GTC TCG TGG GCT CGG AGA TGT GTA TAA GAG ACA GGG ACT ACH VGG GTW TCT AAT) targeting the V4 regions of the 16S rRNA gene (Edwards *et al*., 2015). The amplified products were purified using agarose gel electrophoresis and the Fast Gene Gel/PCR Extraction kit. Paired-end sequencing was performed on an Illumina Miseq sequencer using the Illumina Miseq reagent Kit V3. The obtained sequence data were analyzed at Bioengineering Lab. Co., Japan. They extracted only the read sequences that perfectly matched the primer sequences used for the 5’ end reading of the read sequences obtained using the fastx_barcode_splitter tool of FASTX-Toolkit ver 0.0.14 (https://github.com/agordon/fastx_toolkit). The primer sequences were removed from the extracted reads using fastx_trimer of the FASTX Toolkit. Subsequently, sickle ver 1.3 (https://github.com/najoshi/sickle) was used to discard sequences with a quality value of less than 20 and sequences and their paired sequences that were less than 130 bases in length. The paired-end reads were merged using the FLASH ver. 1.2.11 (Magoč and Salzberg, 2011) merging script under the following conditions: merged sequence length (bp) 250, read length (bp) 230, and overlap length (bp) 10. After removing chimeric and noisy sequences with the dada2 plugin of Qiime 2 ver. 2022.8 (Bolyen *et al*., 2019), representative sequences and ASV tables were generated. The obtained representative sequences were compared with the 97% OTU of the Greengene database (ver. 13_8) using the feature classifier plugin, and taxonomic assignments were made.

### Community analysis based on the 16S rRNA gene

To identify taxa with significantly different relative abundance between treatments, we conducted statistical association analysis using the linear discriminant analysis (LDA) effect size (LEfSe) method (Segata *et al*., 2011). To investigate the biological significance, we subsequently employed the Wilcoxon rank-sum test. Additionally, LDA was employed to estimate the effect size of each differentially abundant feature. Species diversity differences were assessed using Shannon’s index, and community structure differences were assessed using Principal Coordinate Analysis (PCoA) with Bray-Curtis implemented in the *vegan* R package v.4.1.2, both based on OTU counts.

### Statistical analysis

All statistical analyses were performed using EZR version 1.55 (Saitama Medical Center, available at https://www.jichi.ac.jp/saitama-sct/SaitamaHP.files/statmed.html), which is a graphical user interface for R (The R Foundation for Statistical Computing, version 4.1.2) (Kanda, 2013).

### Data availability

The complete genome sequences of *Leucobacter* sp. H87-6, *A. faecalis* H87-33, and *S. maltophilia* H87-40 have been deposited in DDBJ under accession numbers DRA017064, DRA017065, and DRA017066, respectively. The 16S rRNA gene sequences of the bacterial community in plants treated with H87-SC5 or H87-SC3 have been deposited in DDBJ under accession number DRA017018.

## ACKNOWLEDGMENTS

We would like to thank Y. Munetomo for technical assistance and A. Adachi for assisting in data analysis, and M. Umeda, T. Tohge, M. Shimizu, K. Hiruma, H. Adachi, C. Tateda, M. Fuji, S. Yasuda and A. Shinmyo for their helpful comments and suggestions.

## Author contributions

ET, MF, and YS conceived, organized, and supervised the project. ET and YS wrote the manuscript with inputs from the other authors. ET, YT, SK, RM, and DU conducted the experiments and analyzed the data. TM conducted comparative genome analysis on bacteria.

## Supplementary data

**Supplementary Figure 1.** SynComs excluding one strain from H87-SC6 and individual strains do not suppress bacterial wilt disease in tomato seedlings grown in pots with potting soil mix. A) SynComs excluding one strain from H87-SC6 do not show disease-suppressive effects in pot experiments. B) Individual strains that construct H87-SC6 do not significantly reduce the disease index. A) and B) The disease index shows: 0 - no disease symptoms, 1 - up to 25% leaf wilted, 2 - up to 50% leaf wilted, 3 - up to 75% leaf wilted, 4 - up to 100% leaf wilted. The number of plants investigated was 8. A) Experiments were repeated three times. B) Experiments were repeated twice. Data were subjected to one-way ANOVA with Dunnett’s multiple comparison test.

**Supplementary Figure 2.** Disease-suppressive effect of H87-SC6 was high when H87-SC6 colonized tomato plants at a level of 10^5^-10^6^ cfu/g. A) Validation of the relationship between the population of SynCom and disease incidence in a seedling bioassay under sterile conditions. Two test tubes were prepared for each treatment, one was used to evaluate disease symptoms 12 days post inoculation of *R. pseudosolanacearum*, and the other was used to measure the population of SynCom 9 days post seeding. Mixed samples of three tomato plants were used to measure the population of SynCom. B) The population of SynCom in the plant 9 days post seeding affects disease incidence 12 days post inoculation of *R. pseudosolanacearum*. The middle line shows the average disease incidence (The number of plants investigated was 8-10 per tube). The colors of the marks represent different experiments.

**Supplementary Figure 3.** SynCom H87-SC5, which was constructed with five strains, showed a higher disease-suppressive effect than H87-SC6 on tomato seedlings grown in test-tubes with sterile vermiculite. A) Validation of the disease-suppressive effects of SynCom H87-SC6 and SynComs constructed with five strains that make up H87-SC6 in a seedling bioassay under sterile conditions. B) Disease incidence in H87-SC5 treatments (constructed with one *Leucobacter* sp., one *Alcaligenes* sp., and three *Stenotrophomonas* sp.) was significantly lower than in other SynComs 12 days post inoculation of *R. pseudosolanacearum*. Error bars represent the standard deviation of the mean. The middle line shows the average disease incidence (The number of plants investigated was 8-10 per tube. The number of tubes was thirteen for mock controls and H87-SC6 treatment, and three for other treatments.). Data were subjected to one-way ANOVA with Tukey- Kramer’s multiple comparison test. Different letters indicate significant differences at p < 0.05. The colors of the marks represent different experiments.

**Supplementary Figure 4.** Taxa at the genus level with significantly different relative abundance detected by LEfSe. A) Taxa with significantly different relative abundance in plants between H87-SC5 treatment and mock treatment (neither was inoculated with *R. pseudosolanacearum*). B) Taxa with significantly different relative abundance in plants in H87-SC5 treatment between inoculated with *R. pseudosolanacearum* and not inoculated with it. C) Taxa with significantly different relative abundance in plants in treatments between H87-SC5 and H87-SC3 (neither was inoculated with *R. pseudosolanacearum*). D) Taxa with significantly different relative abundance in plants in mock treatment between inoculated and not inoculated with *R. pseudosolanacearum*. Circles indicate the taxa to which the strains constructing H87-SC5 or H87-SC3 belong.

**Supplementary Figure 5.** Comparison of the plant-associated bacterial community structure. A) Principal coordinate analysis (PCoA) showing that β diversity of the plant-associated bacterial community structure is affected by SynCom treatments. B) Shannon indexes showing that α diversity of the plant-associated bacterial community structure among mock treatment, H87-SC5 treatment, and H87-SC3 treatment (no inoculation of *R. pseudosolanacearum*) are not significantly different. Error bars represent the standard deviation of the mean. The middle line shows the average disease incidence (The number of samples was 6 per treatment). Different letters indicate significant differences at p < 0.05.

**Supplementary Figure 6.** SynCom H87-SC6 and H87-SC5 did not inhibit tomato plant growth. A) Validation of non-inhibition of plant growth by SynCom H87-SC6 and H87-SC5 in pot and field experiments. B) The heights of plants treated with H87-SC6 or H87-SC5 are not significantly different from those in mock controls in pot experiments. C) The heights of plants treated with H87-SC6 or H87-SC5 are not significantly different from those in mock controls in field experiments. B) and C) Error bars represent the standard deviation of the mean (The number of plants investigated was 4 per treatment). Data were subjected to one-way ANOVA with Dunnett’s multiple comparison test.

**Supplementary Table 1.** Total of 1,934 tomato plant-associated microbial complexes were collected. Tomato root samples were collected from nine pesticide-free farms, including three farms in Osaka, one farm in Shiga, one farm in Nara, and four farms in Okayama. The collected root samples yielded 1,934 tomato plant-associated microbial complexes.

**Supplementary Table 2.** Average Nucleotide Identity (ANI) between the genomes of the selected strains and representative strains of the same species.

**Supplementary Table 3.** Strain T7-34 exhibited negative disease-suppressive effects against *R. pseudosolanacearum* in both non-sterile and sterile seedling bioassays. The disease-suppressive effects of strain T7-34 were investigated using non-sterile and sterile seedling bioassays, with 8-10 plants being investigated. A negative value in the reduction of disease incidence indicates a negative disease suppressive effect. Taxonomic information for T7-34 was obtained through sequence analysis of the 16S rRNA gene and a BLAST search.

**Supplementary Table 4.** Genome information for H87-6, H87-33, and H87-40.

**Supplementary Table 5.** Gene annotation for disease control-related functions of H87-40, H87-6, and H87-33.

**Supplementary Table 6.** Endophytic bacterial community of plants grown in soil from which the selected strains were isolated. Taxonomic information was obtained through 16S rRNA amplicon sequencing.

**Supplementary Table 7.** Five types of media were used to isolate plant-associated microbial complexes. NYGB, TSB and PDA are nutrient-rich media, on the other hand, R2A and WA are nutrient-poor media.

## Notes

### Competing Interest Statement

The authors have declared no competing interest.

## LITERATURE CITED

Achari, G. A. and Ramesh, R. (2015) Characterization of bacteria degrading 3-hydroxy palmitic acid methyl ester (3OH-PAME), a quorum sensing molecule of *Ralstonia solanacearum*. Letters in applied microbiology, 60(5), 447–455.

Asnicar, F., Thomas, A. M., Beghini, F., Mengoni, C., Manara, S., Manghi, P., Zhu, Q., Bolzan, M., Cumbo, F., May, U., Sanders J. G., Zolfo, M., Kopylova, E., Pasolli, E., Knight, R., Mirarab, S., Huttenhower, C., and Segata, N. (2020). Precise phylogenetic analysis of microbial isolates and genomes from metagenomes using PhyloPhlAn 3.0. Nature communications, 11(1), 2500.

Banerjee, S., Schlaeppi, K., van der Heijden MGA. (2018) Keystone taxa as drivers of microbiome structure and functioning. Nature Reviews Microbiology16: 567–576.

Bates, K. A., and King, K. C. (2021) Leucobacter. Trends in microbiology.

Bayer-Santos, E., Cenens, W., Matsuyama, B. Y., Oka, G. U., Di Sessa, G., Mininel, I. D. V., Alves, T., L., and Farah, C. S. (2019). The opportunistic pathogen *Stenotrophomonas maltophilia* utilizes a type IV secretion system for interbacterial killing. PLoS pathogens, 15(9), e1007651.

Berihu, M., Somera, T. S., Malik, A., Medina, S., Piombo, E., Tal, O., Cohen, M., Ginatt, A., Ofek-Lalzar M., Doron-Faigenboim A., Mazzola M. and Freilich, S. (2023). A framework for the targeted recruitment of crop-beneficial soil taxa based on network analysis of metagenomics data. Microbiome, 11(1), 8.

Bolyen, E., Rideout, J. R., Dillon, M. R., Bokulich, N. A., Abnet, C. C., Al-Ghalith, G. A., Alexander H, Alm EJ, Arumugam M, Asnicar F, et al. (2019). Reproducible, interactive, scalable and extensible microbiome data science using QIIME 2. Nature biotechnology, 37(8), 852–857.

Camargo, A. P., de Souza, R. S., Jose, J., Gerhardt, I. R., Dante, R. A., Mukherjee, S., Kyrpides, N. C., Carazzolle, M. F., Arruda, P. (2023). Plant microbiomes harbor potential to promote nutrient turnover in impoverished substrates of a Brazilian biodiversity hotspot. The ISME journal, 17(3), 354–370.

Compant, S., Samad, A., Faist, H., and Sessitsch, A. (2019) A review on the plant microbiome: ecology, functions, and emerging trends in microbial application. Journal of advanced research, 19, 29–37.

De Vrieze, M., Germanier, F., Vuille, N., and Weisskopf, L. (2018) Combining different potato- associated *Pseudomonas* strains for improved biocontrol of *Phytophthora* infestans. Frontiers in microbiology, 9, 2573.

Edwards, J., Santos-Medellín, C., and Sundaresan, V. (2018) Extraction and 16S rRNA sequence analysis of microbiomes associated with rice roots. Bio-protocol, 8(12), e2884–e2884.

Elhalag, K. M., Messiha, N. A. S., Emara, H. M., and Abdallah, S. A. (2016) Evaluation of antibacterial activity of *Stenotrophomonas maltophilia* against *Ralstonia solanacearum* under different application conditions. Journal of Applied microbiology, 120(6), 1629–1645.

Fatima, T., Mishra, I., Verma, R., and Arora, N. K. (2020) Mechanisms of halotolerant plant growth promoting *Alcaligenes* sp. involved in salt tolerance and enhancement of the growth of rice under salinity stress. 3 Biotech, 10(8), 1-12.

Feil, H., Feil, W. S., Chain, P., Larimer, F., DiBartolo, G., Copeland, A., Lykidis, A., Trong, S., Nolan, M., Goltsman, E., Thiel, J., Malfatti, S., Loper, J. E., Lapidus, A., Detter, J. C., Land, M., Richardson, P. M., Kyrpides, N.C., Ivanova, N., and Lindow, S. E. (2005) Comparison of the complete genome sequences of *Pseudomonas syringae pv. syringae* B728a and *pv. tomato* DC3000. Proceedings of the National Academy of Sciences, 102(31), 11064–11069.

Food and Agriculture Organization of the United Nations (2017) Plant health and food security, http://www.fao.org/3/a-i7829e.pdf.

French, E.B., Gutarra, L., Aley, P., and Elphinstone, J. (1995) Culture media for *Ralstonia solanacearum* isolation, identification and maintenance. Fitopatología 30: 126–130.

Fu, H. Z., Marian, M., Enomoto, T., Hieno, A., Ina, H., Suga, H., and Shimizu, M. (2020) Biocontrol of tomato bacterial wilt by foliar spray application of a novel strain of endophytic *Bacillus* sp. Microbes and environments, 35(4), ME20078.

Gu, S., Wei, Z., Shao, Z., Friman, V. P., Cao, K., Kramer, Y. J., Wang, X., Li, M., Mei, X., Xu, Y., Shen, Q., Kümmerli, R. and Jousset, A. (2020) Competition for iron drives phytopathogen control by natural rhizosphere microbiomes. Nature Microbiology, 5(8), 1002–1010.

Hardoim, P. R., van Overbeek, L. S., and van Elsas, J. D. (2008) Properties of bacterial endophytes and their proposed role in plant growth. Trends in microbiology, 16(10), 463–471.

Hardoim, P. R., van Overbeek, L. S., Berg, G., Pirttilä, A. M., Compant, S., Campisano, A., Döring M. and Sessitsch, A. (2015). The hidden world within plants: ecological and evolutionary considerations for defining functioning of microbial endophytes. Microbiology and molecular biology reviews, 79(3), 293–320.

Hikichi, Y., Mori, Y., Ishikawa, S., Hayashi, K., Ohnishi, K., Kiba, A. and Kai, K. (2017) Regulation involved in colonization of intercellular spaces of host plants in *Ralstonia solanacearum*. Front. Plant Sci. 8, 967.

Jain, C., Rodriguez-R, L. M., Phillippy, A. M., Konstantinidis, K. T., and Aluru, S. (2018) High throughput ANI analysis of 90K prokaryotic genomes reveals clear species boundaries. Nature Communications 9, 5114.

Kanda, Y. (2013). Investigation of the freely available easy-to-use software ‘EZR’for medical statistics. Bone marrow transplantation, 48(3), 452–458.

Kolmogorov, M., Yuan, J., Lin, Y., and Pevzner, P. A. (2019). Assembly of long, error-prone reads using repeat graphs. Nature biotechnology, 37(5), 540–546.

Kozlov, A. M., Darriba, D., Flouri, T., Morel, B. and Stamatakis, (2019) A. RAxML-NG: a fast, scalable and user-friendly tool for maximum likelihood phylogenetic inference. Bioinformatics 35, 4453–4455.

Kelman, A. (1954) The relationship of pathogenicity of *Pseudomonas solanacearum* to colony appearance in a tetrazolium medium. Phytopathology, 44(12).

Kumatani, T., Yoshimi, Y., Nakayashiki, and H., Aino, M. (2009) Phylogenetic analyses of plant- growth-promoting rhizobacteria isolated from tomato, lettuce, and Japanese pepper plants in Hyogo Prefecture, Japan, Journal of General Plant Pathology, 75, 316–321.

Kwak, M. J., Kong, H. G., Choi, K., Kwon, S. K., Song, J. Y., Lee, J., Lee, P. A., Choi, S. Y., Seo, M., Lee, H. J., Jung, E. J., Park, H., Roy, N., Kim, H., Lee, M. M., Rubin, E. M., Lee, S. W.,and Kim, J. F. (2018) Rhizosphere microbiome structure alters to enable wilt resistance in tomato. Nature Biotechnology, 36,1100–1117.

Magoč, T., and Salzberg, S. L. (2011). FLASH: fast length adjustment of short reads to improve genome assemblies. Bioinformatics, 27(21), 2957–2963.

Mansfield, J., Genin, S., Magori, S., Citovsky, V., Sriariyanum, M., Ronald, Ronald, P., Dow, M., Verdier, V., Beer, S. V., Machado, M. A., Toth, I., GEORGE Salmond, G., and Foster, G. D. (2012) Top 10 plant pathogenic bacteria in molecular plant pathology. Molecular plant pathology, 13(6), 614–629.

Marian, M., Nishioka, T., Koyama, H., Suga, H., and Shimizu, M. (2018) Biocontrol potential of *Ralstonia* sp. TCR112 and *Mitsuaria* sp. TWR114 against tomato bacterial wilt. Applied Soil Ecology, 128, 71-80.

Mazzola, M., and Freilich, S. (2017) Prospects for biological soilborne disease control: application of indigenous versus synthetic microbiomes. Phytopathology 107, 256–263.

Messiha, N. A. S., Van Diepeningen, A. D., Farag, N. S., Abdallah, S. A., Janse, J. D., and Van Bruggen, A. H. C. (2007) *Stenotrophomonas maltophilia*: a new potential biocontrol agent of *Ralstonia solanacearum*, causal agent of potato brown rot. European journal of plant pathology, 118(3), 211–225.

Nas, M. Y., White, R. C., DuMont, A. L., Lopez, A. E., and Cianciotto, N. P. (2019) *Stenotrophomonas maltophilia* encodes a VirB/VirD4 type IV secretion system that modulates apoptosis in human cells and promotes competition against heterologous bacteria, including *Pseudomonas aeruginosa*. Infection and immunity, 87(9), 10–1128.

Niu, B., Paulson, J. N., Zheng, X., and Kolter, R. (2017). Simplified and representative bacterial community of maize roots. Proceedings of the National Academy of Sciences, 114(12), E2450–E2459.

Parks, D. H., Imelfort, M., Skennerton, C. T., Hugenholtz, P., and Tyson, G. W. (2015). CheckM: assessing the quality of microbial genomes recovered from isolates, single cells, and metagenomes. Genome research, 25(7), 1043–1055.

Phae, C. G., Shoda, M., Kita, N., Nakano, M., and Ushiyama, K. (1992) Biological control of crown and root rot and bacterial wilt of tomato by *Bacillus subtilis* NB22. Japanese Journal of Phytopathology, 58(3), 329–339.

Prior, P., Ailloud, F., Dalsing, B. L., Remenant, B., Sanchez, B., and Allen, C. (2016) Genomic and proteomic evidence supporting the division of the plant pathogen *Ralstonia solanacearum* into three species. BMC genomics, 17(1), 1–11.

Reinhold-Hurek, B., Bünger, W., Burbano, C.S., Sabale, M., and Hurek, T. (2015) Roots shaping their microbiome: global hotspots for microbial activity, Annu Rev Phytopathol, 53, 403–424.

Roberts, D. P., Denny, T. P., and Schell, M. A. (1988) Cloning of the egl gene of *Pseudomonas solanacearum* and analysis of its role in phytopathogenicity. Journal of Bacteriology, 170(4), 1445–1451.

Schell, M. A. (2000) Control of virulence and pathogenicity genes of *Ralstonia Solanacearum* by an elaborate sensory network, Annu. Rev. Phytopathol. 38:263–92.

Seemann, T. (2014). Prokka: rapid prokaryotic genome annotation. Bioinformatics, 30(14), 2068–2069.

Segata, N., Izard, J., Waldron, L., Gevers, D., Miropolsky, L., Garrett, W. S., and Huttenhower, C. (2011) Metagenomic biomarker discovery and explanation. Genome biology, 12, 1–18.

Singh, R. P., and Jha, P. N. (2017) The PGPR *Stenotrophomonas maltophilia* SBP-9 augments resistance against biotic and abiotic stress in wheat plants. Frontiers in microbiology, 8, 1945.

Singh, R., Kalra, A., Ravish, B. S., Divya, S., Parameswaran, T. N., Srinivas, K. V. N. S., and Bagyaraj, D. J. (2012) Effect of potential bioinoculants and organic manures on root-rot and wilt, growth, yield and quality of organically grown Coleus forskohlii in a semiarid tropical region of Bangalore (India). Plant pathology, 61(4), 700–708.

Wang, X., Wei, Z., Yang, K., Wang, J., Jousset, A., Xu, Y., Shen, Q. and Friman, V. P. (2019) Phage combination therapies for bacterial wilt disease in tomato. Nature Biotechnology, 37(12), 1513–1520.

Wei, Z., Huang, J., Tan, S., Mei, X., Shen, Q., and Xu, Y. (2013) The congeneric strain *Ralstonia pickettii* QL-A6 of *Ralstonia solanacearum* as an effective biocontrol agent for bacterial wilt of tomato. Biological Control, 65(2), 278–285.

Xue, Q. Y., Chen, Y., Li, S. M., Chen, L. F., Ding, G. C., Guo, D. W., and Guo, J. H. (2009) Evaluation of the strains of *Acinetobacter* and *Enterobacter* as potential biocontrol agents against Ralstonia wilt of tomato. Biological Control, 48(3), 252–258.

Yuan, S., Wang, L., Wu, K., Shi, J., Wang, M., Yang, X., Shen, O. and Shen, B. (2014) Evaluation of *Bacillus*-fortified organic fertilizer for controlling tobacco bacterial wilt in greenhouse and field experiments. Applied soil ecology, 75, 86–94.

Yuliar, Nion, Y. A., and Toyota, K. (2015) Recent trends in control methods for bacterial wilt diseases caused by *Ralstonia solanacearum*. *Microbes and environments*, ME14144.

