## Supplementary figures for "Biocontrol of bacterial wilt disease using plant-associated bacterial communities in tomato"

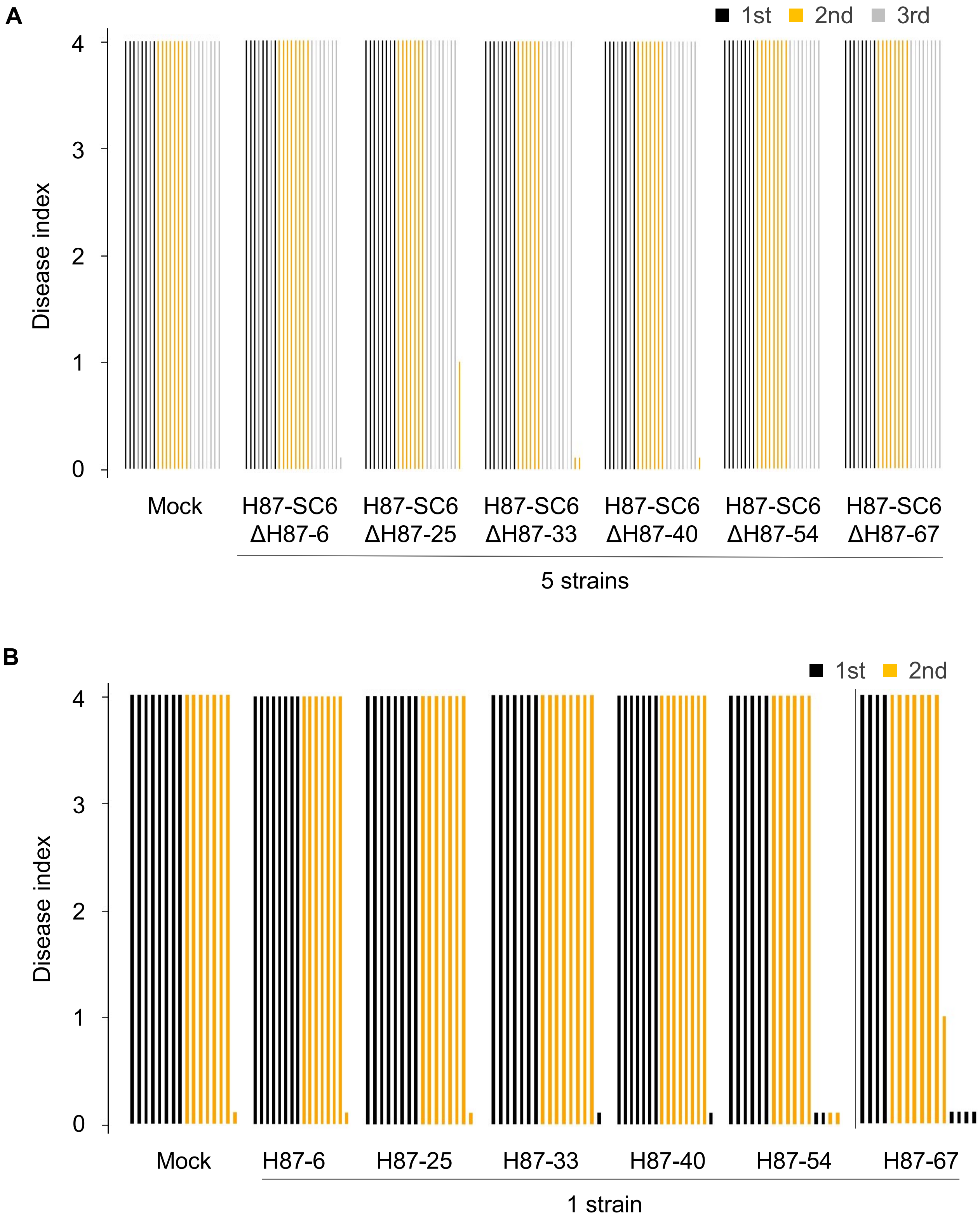

A
1st
2nd
3rd
4
3
2
1
0
Disease index
Mock
H87-SC6 ΔH87-6
H87-SC6 ΔH87-25
H87-SC6 ΔH87-33
H87-SC6 ΔH87-40
H87-SC6 ΔH87-54
H87-SC6 ΔH87-67
5 strains
B
1st
2nd
4
3
2
1
0
Disease index
Mock
H87-6
H87-25
H87-33
H87-40
H87-54
H87-67
1 strain

**Supplementary Figure 1.** SynComs excluding one strain from H87-SC6 and individual strains do not suppress bacterial wilt disease in tomato seedlings grown in pots with potting soil mix. A) SynComs excluding one strain from H87-SC6 do not show disease-suppressive effects in pot experiments. B) Individual strains that construct H87-SC6 do not significantly reduce the disease index. A) and B) The disease index shows: 0 - no disease symptoms, 1 - up to 25% leaf wilted, 2 - up to 50% leaf wilted, 3 - up to 75% leaf wilted, 4 - up to 100% leaf wilted. The number of plants investigated was 8. A) Experiments were repeated three times. B) Experiments were repeated twice. Data were subjected to one-way ANOVA with Dunnett's multiple comparison test.

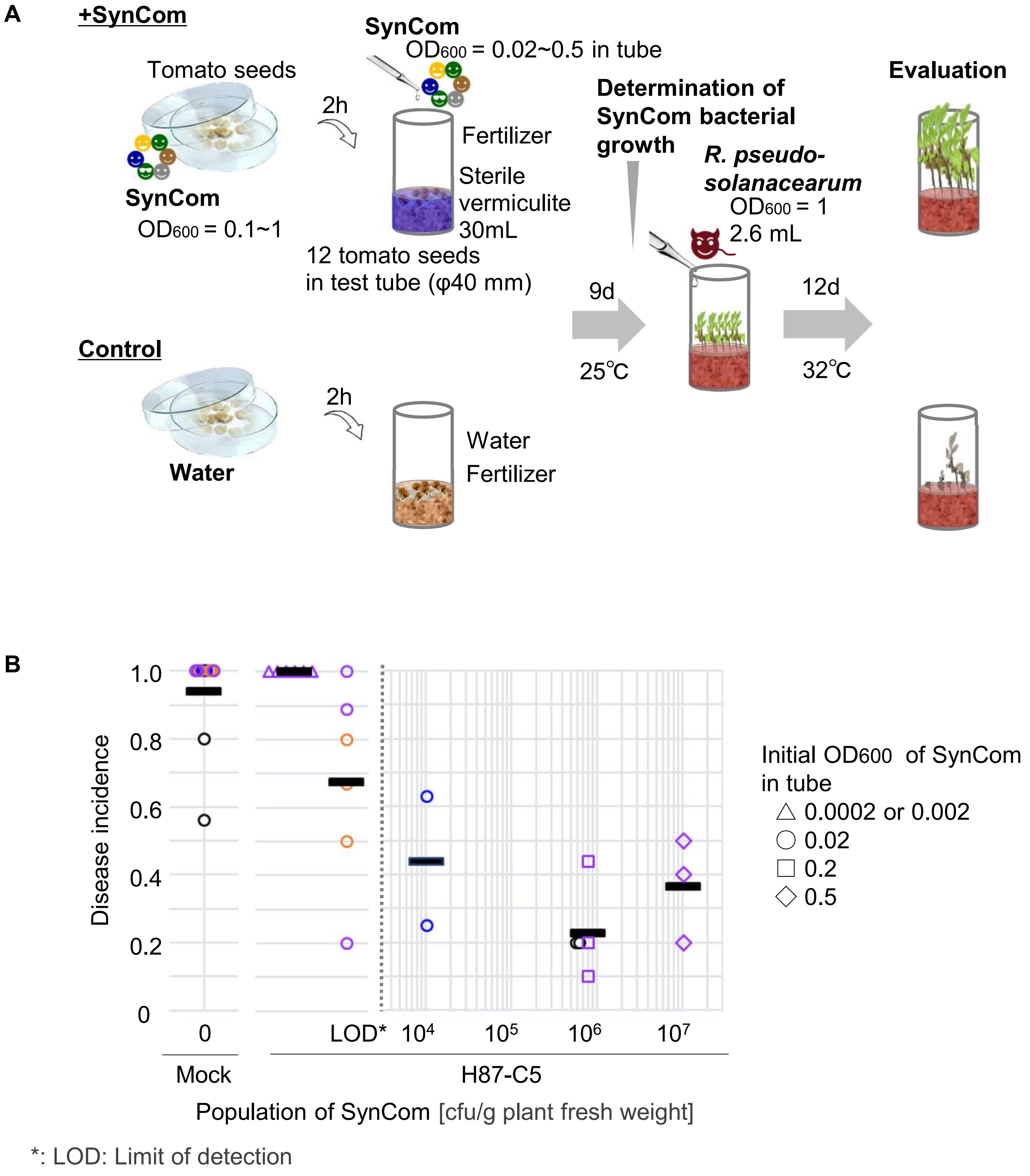


**Supplementary Figure 2.** Disease-suppressive effect of H87-SC6 was high when H87-SC6 colonized tomato plants at a level of $10^5$-$10^6$ cfu/g. A) Validation of the relationship between the population of SynCom and disease incidence in a seedling bioassay under sterile conditions. Two test tubes were prepared for each treatment, one was used to evaluate disease symptoms 12 days post inoculation of *R. pseudosolanacearum*, and the other was used to measure the population of SynCom 9 days post seeding. Mixed samples of three tomato plants were used to measure the population of SynCom. B) The population of SynCom in the plant 9 days post seeding affects disease incidence 12 days post inoculation of *R. pseudosolanacearum*. The middle line shows the average disease incidence (The number of plants investigated was 8-10 per tube). The colors of the marks represent different experiments.

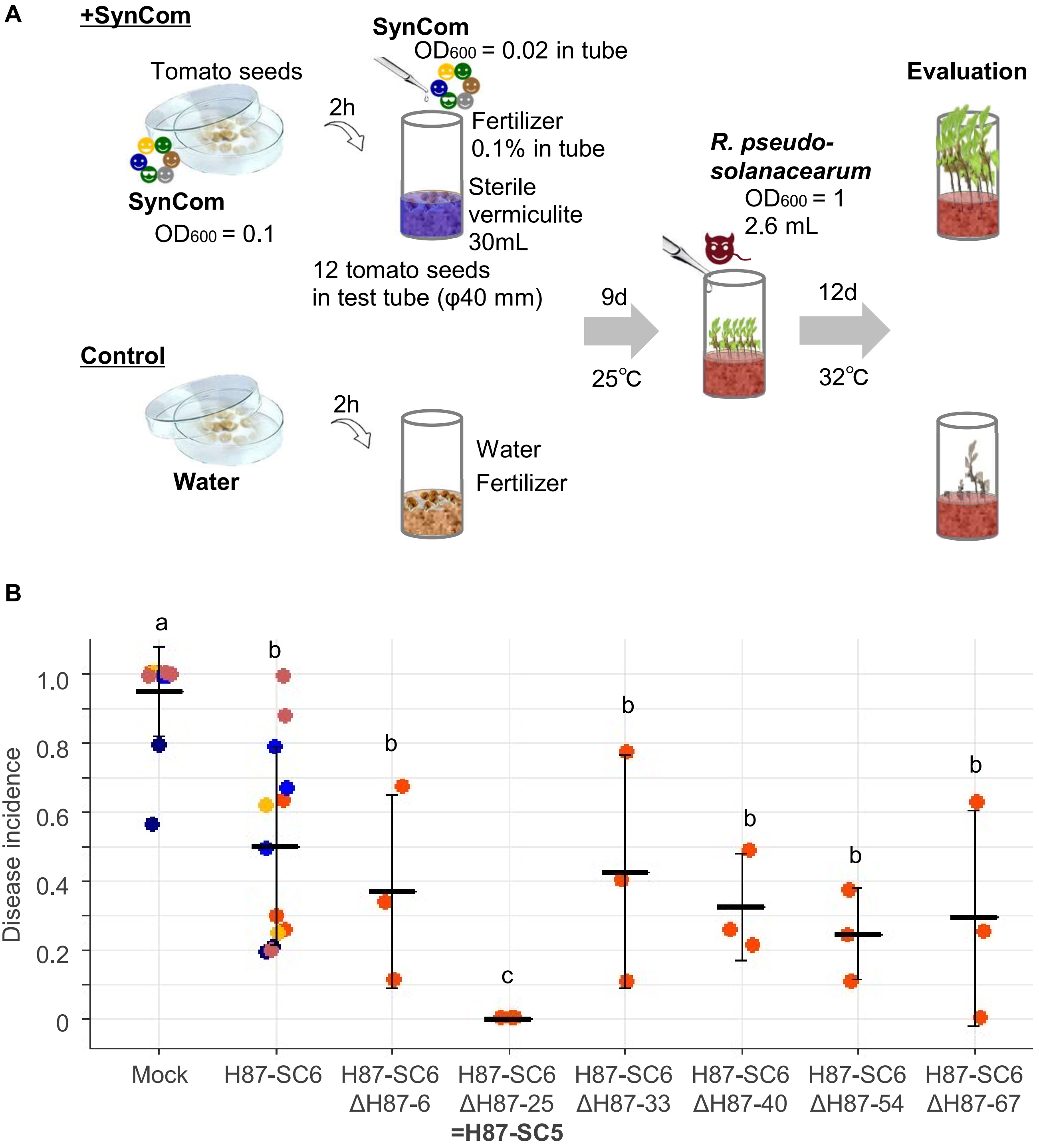


**Supplementary Figure 3.** SynCom H87-SC5, which was constructed with five strains, showed a higher disease-suppressive effect than H87-SC6 on tomato seedlings grown in test-tubes with sterile vermiculite. A) Validation of the disease-suppressive effects of SynCom H87-SC6 and SynComs constructed with five strains that make up H87-SC6 in a seedling bioassay under sterile conditions. B) Disease incidence in H87-SC5 treatments (constructed with one *Leucobacter* sp., one *Alcaligenes* sp., and three *Stenotrophomonas* sp.) was significantly lower than in other SynComs 12 days post inoculation of *R. pseudosolanacearum*. Error bars represent the standard deviation of the mean. The middle line shows the average disease incidence (The number of plants investigated was 8-10 per tube. The number of tubes was thirteen for mock controls and H87-SC6 treatment, and three for other treatments.). Data were subjected to one-way ANOVA with Tukey-Kramer's multiple comparison test. Different letters indicate significant differences at $p < 0.05$. The colors of the marks represent different experiments.

A

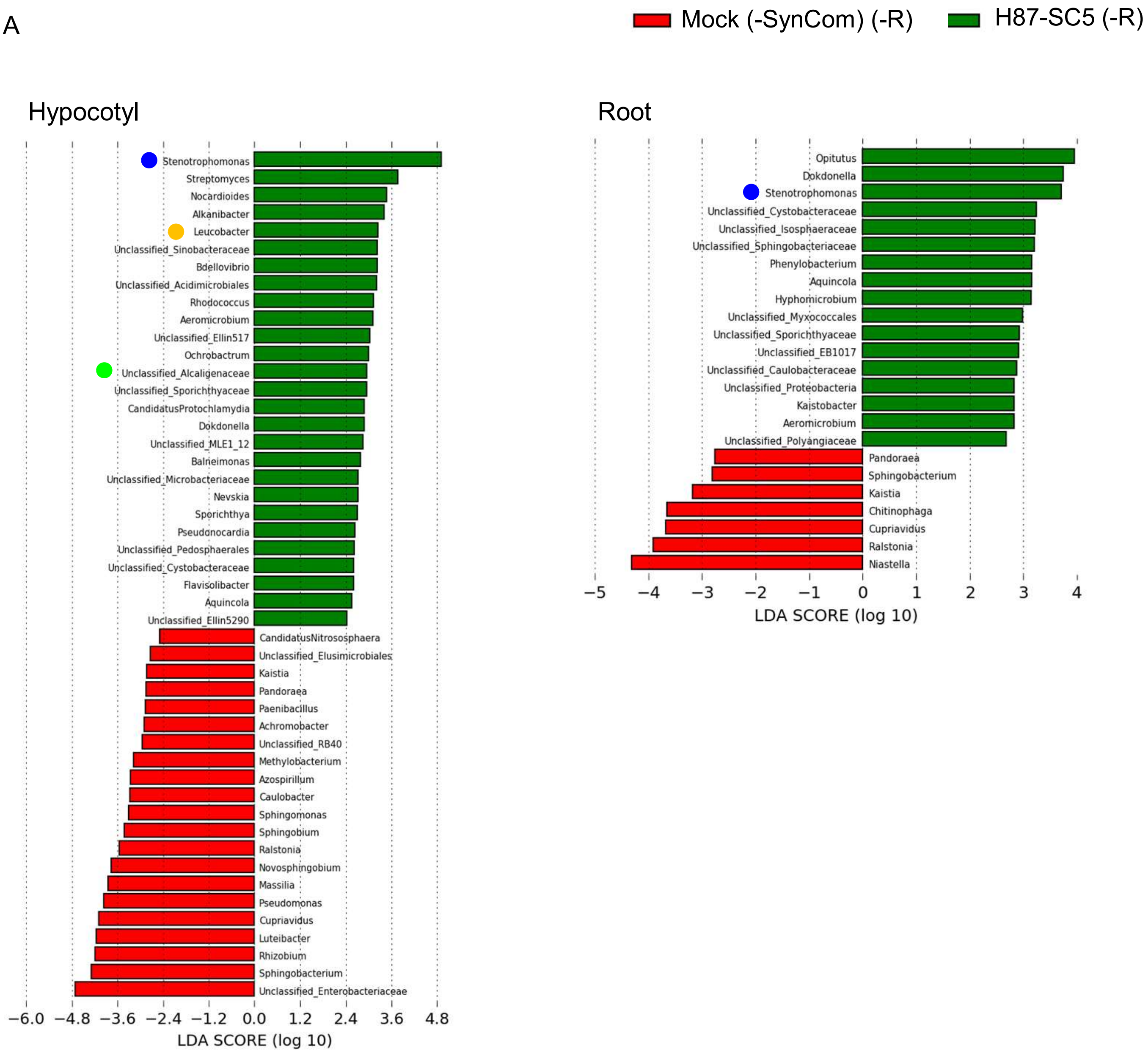


-R: no inoculation of *R. pseudosolanacearum*

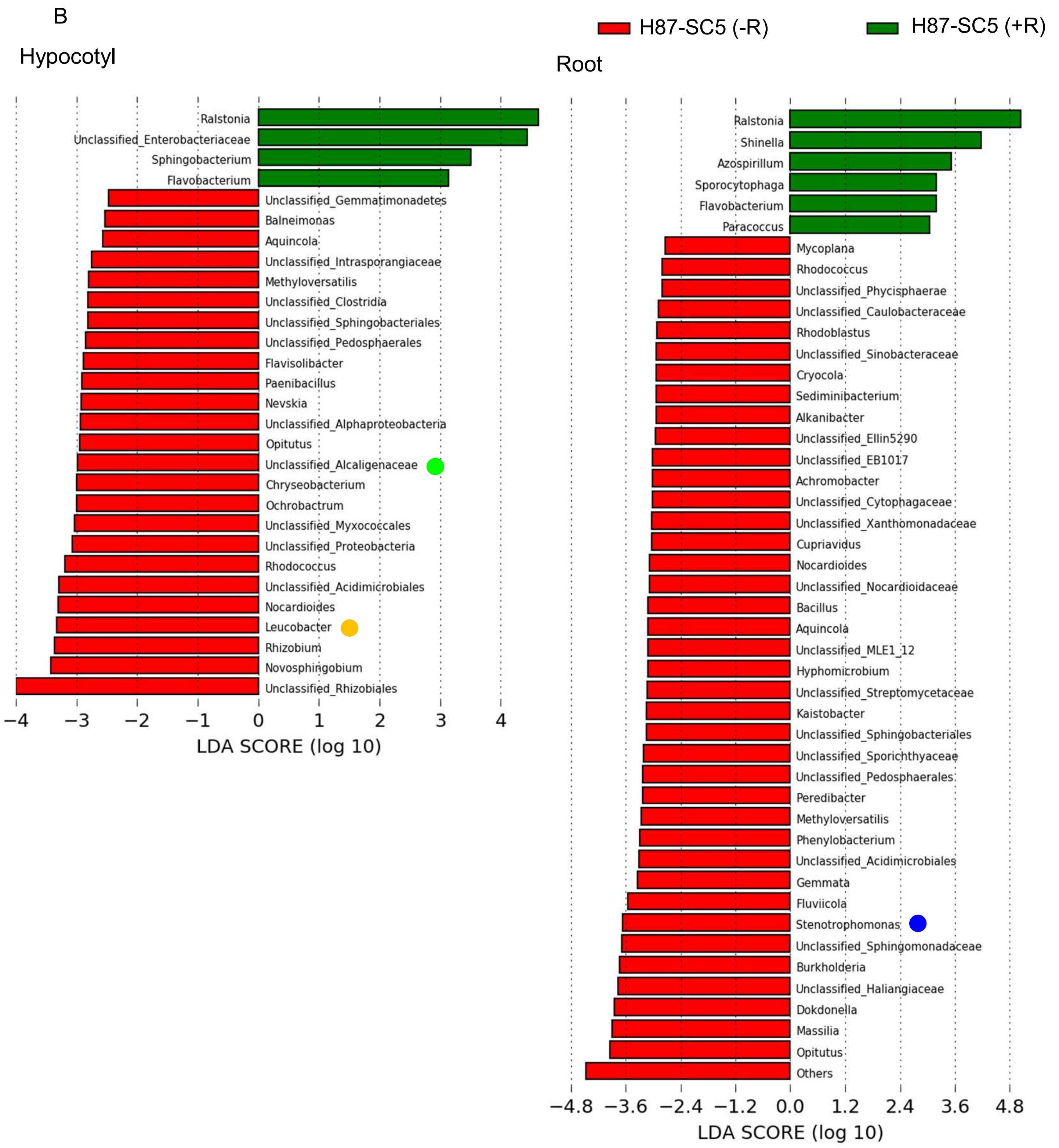


-R: no inoculation of *R. pseudosolanacearum*
+R: inoculation of *R. pseudosolanacearum*

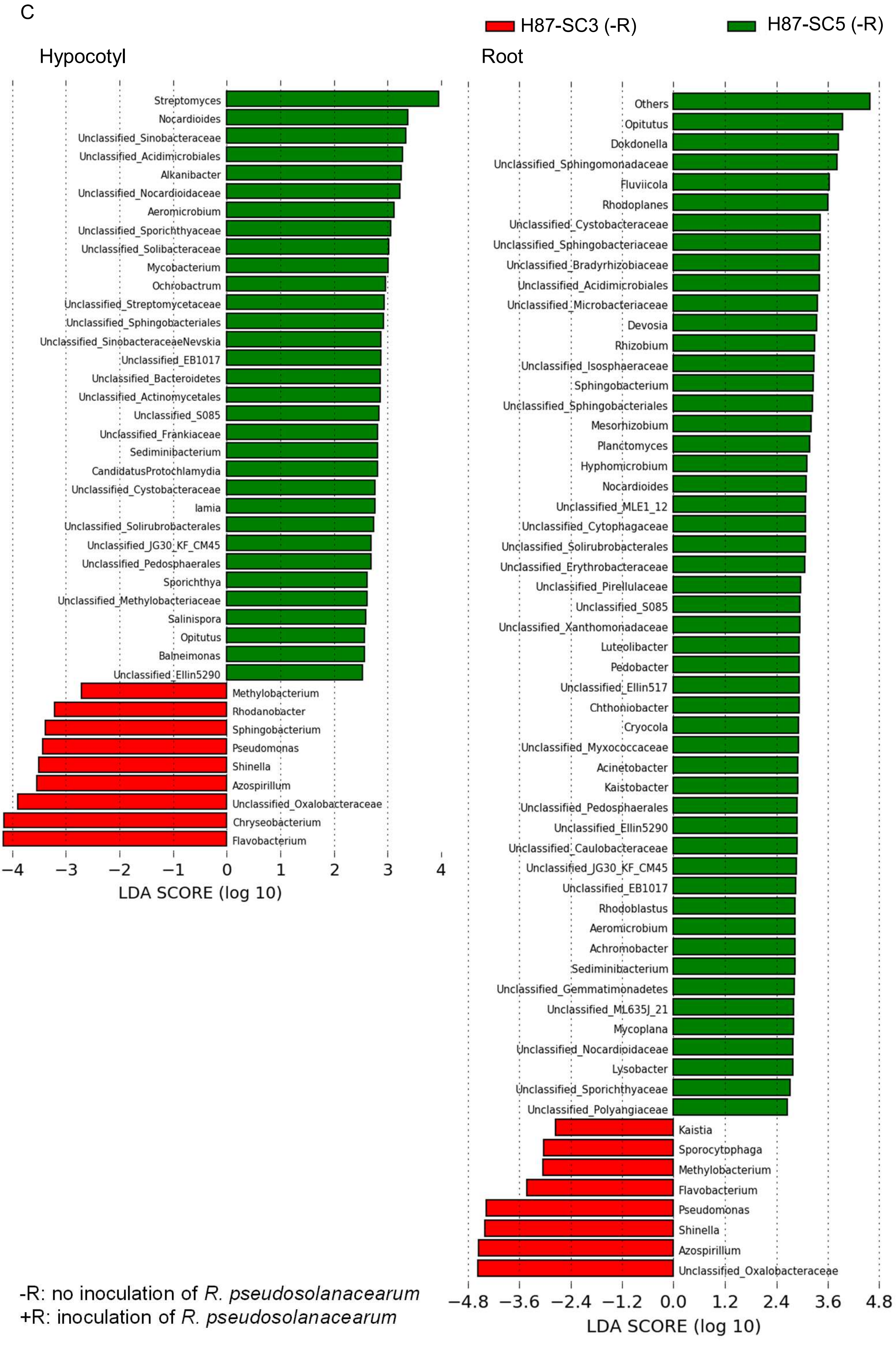


-R: no inoculation of *R. pseudosolanacearum*
+R: inoculation of *R. pseudosolanacearum*

D

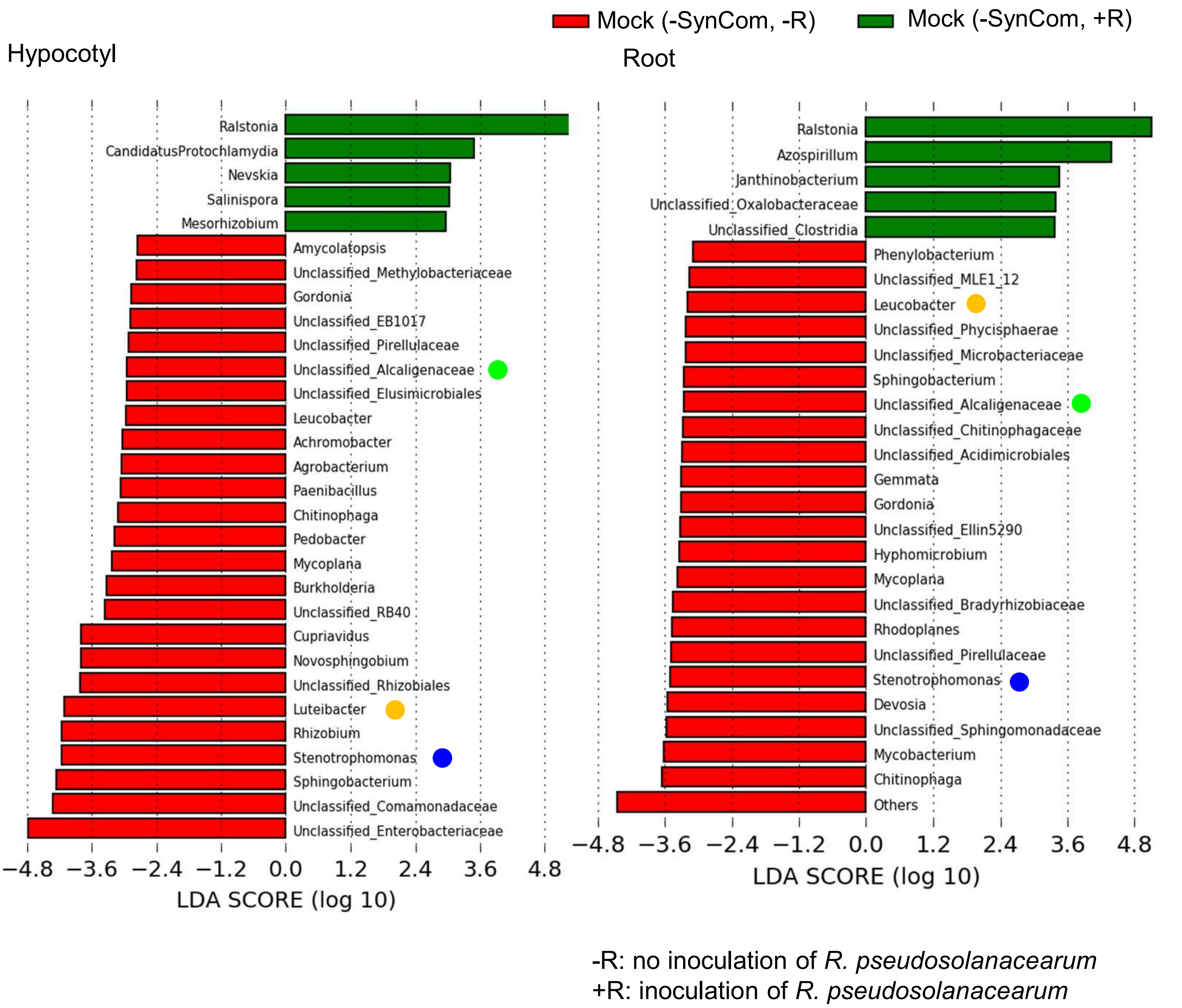


**Supplementary Figure 4.** Taxa at the genus level with significantly different relative abundance detected by LEfSe. A) Taxa with significantly different relative abundance in plants between H87-SC5 treatment and mock treatment (neither was inoculated with *R. pseudosolanacearum*). B) Taxa with significantly different relative abundance in plants in H87-SC5 treatment between inoculated with *R. pseudosolanacearum* and not inoculated with it. C) Taxa with significantly different relative abundance in plants in treatments between H87-SC5 and H87-SC3 (neither was inoculated with *R. pseudosolanacearum*). D) Taxa with significantly different relative abundance in plants in mock treatment between inoculated and not inoculated with *R. pseudosolanacearum*. Circles indicate the taxa to which the strains constructing H87-SC5 or H87-SC3 belong.

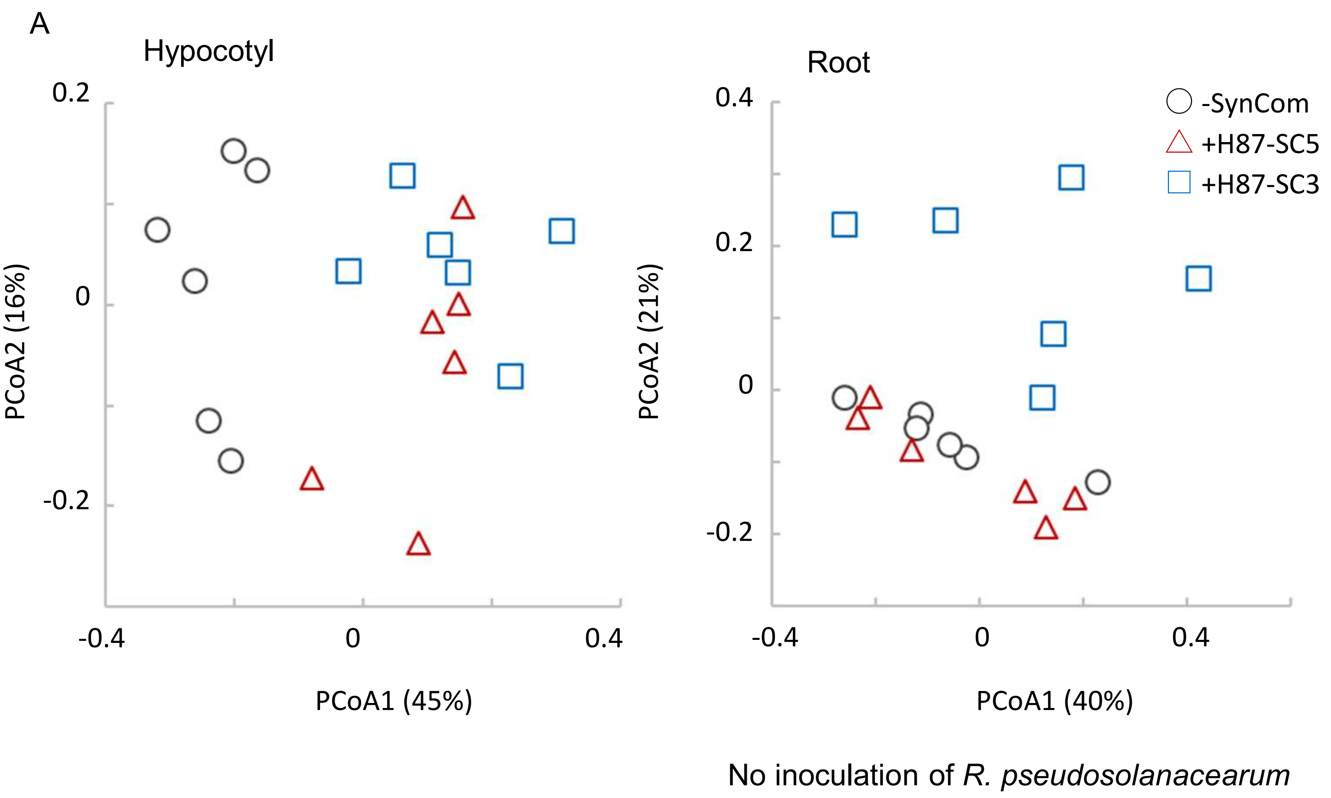


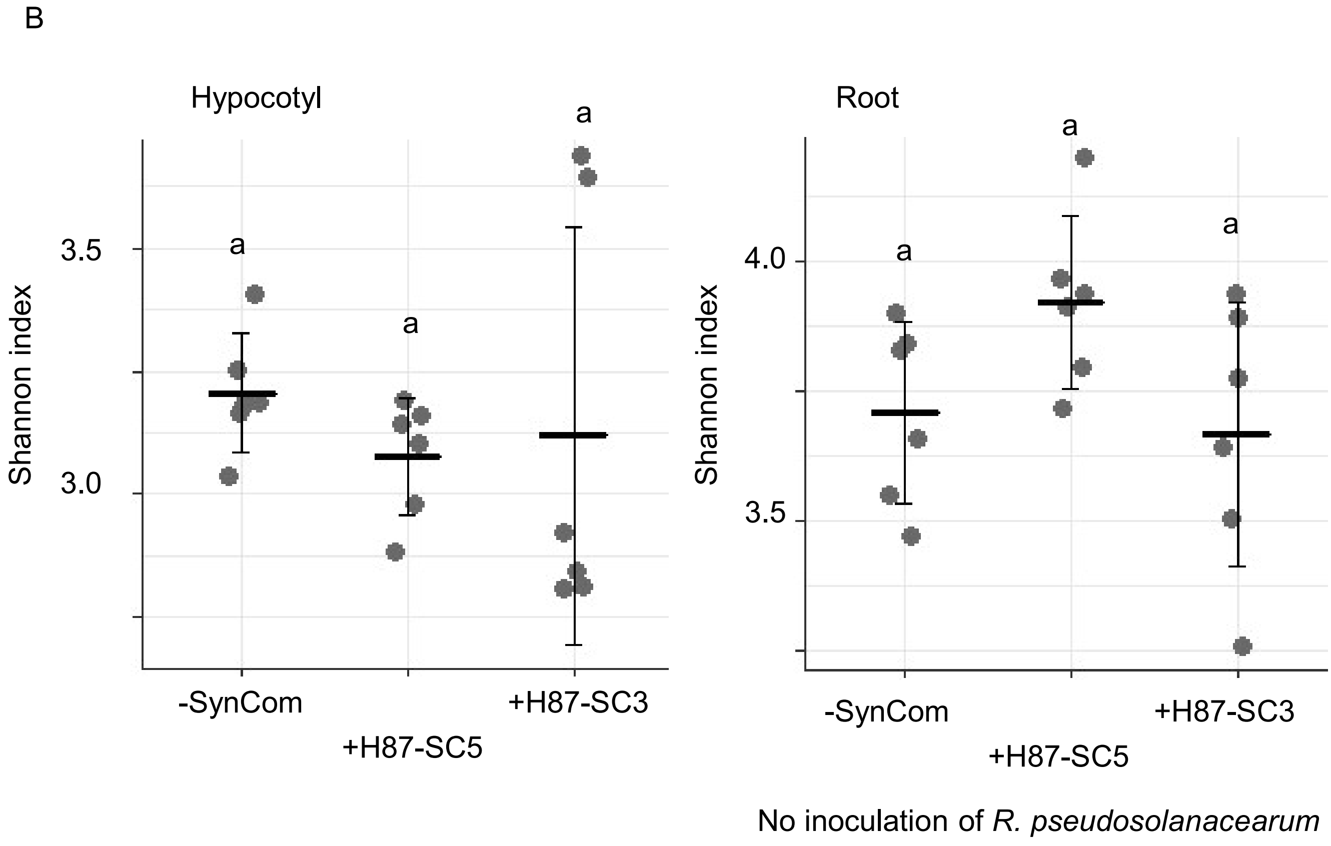


**Supplementary Figure 5.** Comparison of the plant-associated bacterial community structure. A) Principal coordinate analysis (PCoA) showing that β diversity of the plant-associated bacterial community structure is affected by SynCom treatments. B) Shannon indexes showing that α diversity of the plant-associated bacterial community structure among mock treatment, H87-SC5 treatment, and H87-SC3 treatment (no inoculation of *R. pseudosolanacearum*) are not significantly different. Error bars represent the standard deviation of the mean. The middle line shows the average disease incidence (The number of samples was 6 per treatment). Different letters indicate significant differences at $p < 0.05$.

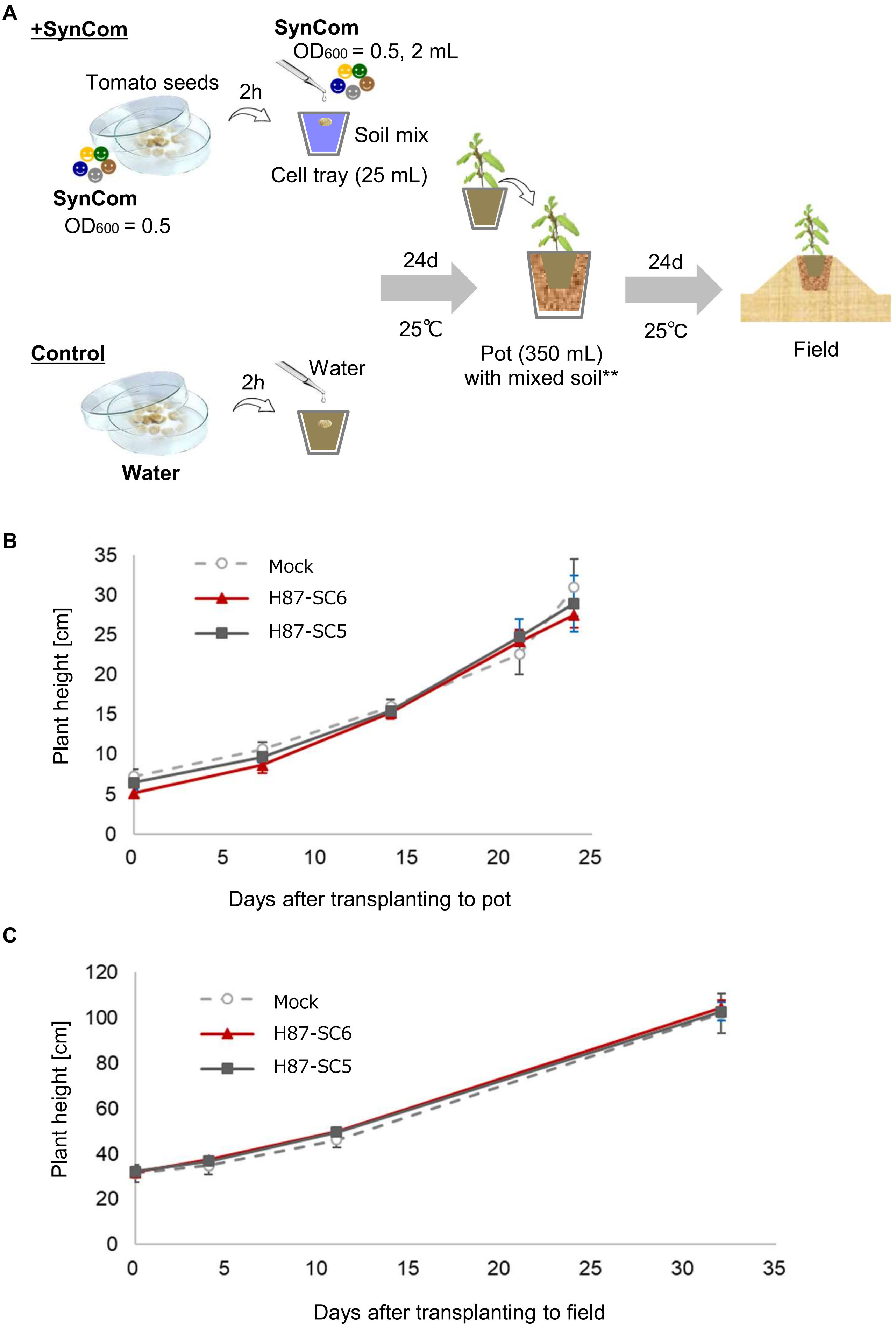
A
+SynCom
Tomato seeds
SynCom
OD600 = 0.5
2h
SynCom
OD600 = 0.5, 2 mL
Soil mix
Cell tray (25 mL)
24d
25℃
Pot (350 mL)
with mixed soil**
24d
25℃
Field
Control
Water
2h
Water
B
Mock
H87-SC6
H87-SC5
Plant height [cm]
Days after transplanting to pot
C
Mock
H87-SC6
H87-SC5
Plant height [cm]
Days after transplanting to field

**Supplementary Figure 6.** SynCom H87-SC6 and H87-SC5 did not inhibit tomato plant growth. A) Validation of non-inhibition of plant growth by SynCom H87-SC6 and H87-SC5 in pot and field experiments. B) The heights of plants treated with H87-SC6 or H87-SC5 are not significantly different from those in mock controls in pot experiments. C) The heights of plants treated with H87-SC6 or H87-SC5 are not significantly different from those in mock controls in field experiments. B) and C) Error bars represent the standard deviation of the mean (The number of plants investigated was 4 per treatment). Data were subjected to one-way ANOVA with Dunnett's multiple comparison test.
