## Supplementary tables for "Biocontrol of bacterial wilt disease using plant-associated bacterial communities in tomato"

Supplementary Table 1. Total of 1,934 tomato plant-associated microbial complexes were collected.

| Field code | Location | Tomato cultivar from which plant-associated microbial complexes were isolated | Number of root samples | Number of microbial complexes |
|---|---|---|---|---|
| Y1 | Osaka, Japan | Frutica (Takii Seed Co., Japan) | 1 | 67 |
| Y2 | | Chika (Takii Seed Co., Japan) | 3 | 308 |
| | | Frutica (Takii Seed Co., Japan) | 1 | 52 |
| | | Momotaro-8 (Takii Seed Co., Japan) | 2 | 163 |
| T | | Unknown | 1 | 79 |
| H | Shiga, Japan | Momotaro-8 (Takii Seed Co., Japan) | 2 | 185 |
| | | Green Save (rootstock, Takii Seed Co., Japan) and Momotaro-8 (scion, Takii Seed Co., Japan) | 1 | 82 |
| N | Nara, Japan | Kyoryoku-beiju (Takii Seed Co., Japan) | 1 | 134 |
| | | Chika (Takii Seed Co., Japan) | 1 | 108 |
| NY1 | Okayama, Japan | Ganbarune-forte (rootstock, Aisan Seed Co.) and Momotaro-8 (scion, Takii Seed Co., Japan)) | 1 | 76 |
| | | Ganbarune-forte (rootstock, Aisan Seed Co.) and Chika (scion, Takii Seed Co., Japan) | 2 | 212 |
| | | Momotaro-8 (Takii Seed Co., Japan) | 2 | 222 |
| NY2 | | Chika (Takii Seed Co., Japan) | 1 | 67 |
| M | | Chika (Takii Seed Co., Japan) | 1 | 85 |
| YK | | Chika (Takii Seed Co., Japan) | 1 | 94 |
| | | | Total | 1,934 |

Tomato root samples were collected from nine pesticide-free farms, including three farms in Osaka, one farm in Shiga, one farm in Nara, and four farms in Okayama. The collected root samples yielded 1,934 tomato plant-associated microbial complexes.

Supplementary Table 2. Average Nucleotide Identity (ANI) between the genomes of the selected strains and representative strains of the same species.

| Query strain | Reference Genome | Reference NCBI accession | ANI | Isolation source |
|---|---|---|---|---|
| H87-6 | *Leucobacter komagatae* DSM 8803$^{T}$ | GCA_006716085.1 | 93.92 | culture contaminant |
| | *Leucobacter aridicollis* DSM 17380$^{T}$ | GCA_013409595.1 | 86.16 | activated sludge |
| H87-33 | *Alcaligenes faecalis* PSM4 | GCA_023702785.1 | 99.59 | landfill soil |
| | *A. faecalis* PSM8 | GCA_023702805.1 | 99.55 | landfill soil |
| | *A. faecalis* AU14 | GCA_005311025.1 | 98.23 | wheat roots |
| | *A. faecalis* D334 | GCA_023101245.1 | 98.22 | mangrove sediment |
| | *A. faecalis* MB207 | GCA_002082085.1 | 98.21 | tannery effluent |
| | *A. faecalis* subsp. *phenolicus* DSM 16503$^{T}$ | GCA_026344105.1 | 96.22 | wastewater |
| H87-40 | *Stenotrophomonas maltophilia* SM-29 | GCA_021554555.1 | 98.78 | human(nosocomial infection) |
| | *S. maltophilia* ZC_2004 | GCA_002738675.1 | 98.77 | human lung |
| | *S. maltophila* S249-2 | GCA_024128235. | 98.75 | chicken |
| | *S. maltophilia* JS56-1 | GCA_013184135.1 | 98.72 | chicken caecum |
| | *S. maltophilia* SMYN45 | GCA_006120775.1 | 98.71 | human sputum |
| | *S. maltophilia* NMG-F-49-2 | GCA_022630995.1 | 98.70 | cattle feces |
| | *S. maltophila* NCTC 10257$^{T}$ | GCA_900186865.1 | 98.11 | oropharyngeal region of human |

T: Type strain

Supplementary Table 3. Strain T7-34 exhibited negative disease-suppressive effects against R. pseudosolanacearum in both non-sterile and sterile seedling bioassays.

| Microbial strain | Reduction of disease incidence [%] | | | | Taxonomy |
|---|---|---|---|---|---|
| | under non-sterile condition | under sterile condition | | | |
| | 1st | 1st | 2nd | 3rd | |
| T7-34 | -10 | -60 | -62 | -40 | *Sphingomonas koreensis* |

The disease-suppressive effects of strain T7-34 were investigated using non-sterile and sterile seedling bioassays, with 8-10 plants being investigated. A negative value in the reduction of disease incidence indicates a negative disease suppressive effect. Taxonomic information for T7-34 was obtained through sequence analysis of the 16S rRNA gene and a BLAST search.

Supplementary Table 4. Genome information for H87-6, H87-33 and H87-40.

| | Genome information | | |
|---|---|---|---|
| | H87-6 | H87-33 | H87-40 |
| PacBio sequencing | | | |
| Average read length [bp] | 12,095 | 16,254 | 14,222 |
| Sequencing throughput [bp] | 2.16E+08 | 2.15E+08 | 2.18E+08 |
| Assembly status (Check M) | | | |
| Genome size [bp] | 3.48.E+06 | 4.38.E+06 | 4.56.E+06 |
| Completeness [%] | 99 | 100 | 100 |
| GC content [%] | 66.2 | 56.4 | 66.5 |
| Annotation | | | |
| Taxonomy | *Leucobacter* sp. | *Alcaligenes faecalis* | *Stenotrophomonas maltophilia* |
| Number of CDS | 3,101 | 4,019 | 4,050 |

CDS: Protein coding sequence

Supplementary Table 5. Gene annotation for disease control-related functions of H87-40, H87-6, and H87-33.

| Category | Gene locus | Gene annotation (Prokka) | Gene annotation (KEGG) | KEGG Orthology ID | Reference gene locus* | Protein sequence identity between H87-40-b and Reference gene locus* | Additional Information† |
|---|---|---|---|---|---|---|---|
| T4SS | H87-40_01368 | hypothetical protein | type IV secretion system protein VirB6 | K03201 | Smlt2997 | 100.0 | - |
| T4SS | H87-40_01369 | hypothetical protein | #N/A | #N/A | Smlt2998 | 100.0 | - |
| T4SS | H87-40_01370 | Type IV secretion system protein virB4 | type IV secretion system protein VirB4 [EC:7.4.2.8] | K03199 | Smlt2999 | 99.9 | - |
| T4SS | H87-40_01371 | hypothetical protein | type IV secretion system protein VirB3 | K03198 | Smlt3000 | 100.0 | - |
| T4SS | H87-40_01372 | hypothetical protein | type IV secretion system protein VirB2 | K03197 | Smlt3001 | 100.0 | - |
| T4SS | H87-40_01373 | hypothetical protein | type IV secretion system protein VirB1 | K03194 | Smlt3002 | 99.4 | - |
| T4SS | H87-40_01374 | Type IV secretion system protein VirB11 | type IV secretion system protein VirB11 [EC:7.4.2.8] | K03196 | Smlt3003 | 100.0 | - |
| T4SS | H87-40_01375 | Protein virB10 | type IV secretion system protein VirB10 | K03195 | Smlt3004 | 100.0 | - |
| T4SS | H87-40_01376 | Type IV secretion system protein PtlF | type IV secretion system protein VirB9 | K03204 | Smlt3005 | 100.0 | - |
| T4SS | H87-40_01377 | hypothetical protein | type IV secretion system protein VirB8 | K03203 | Smlt3006 | 100.0 | - |
| T4SS | H87-40_01378 | hypothetical protein | #N/A | #N/A | Smlt3007 | 100.0 | - |
| T4SS | H87-40_01379 | Protein VirD4 | type IV secretion system protein VirD4 [EC:7.4.2.8] | K03205 | Smlt3008 | 100.0 | - |
| Putative T4SS effector | H87-40_01363 | hypothetical protein | #N/A | #N/A | Smlt2990 | 99.8 | lipase |
| Putative T4SS effector | H87-40_01365 | hypothetical protein | putative chitinase | K03791 | Smlt2992 | 99.4 | lysozyme-like hydrolase |
| Putative T4SS effector | H87-40_02644 | hypothetical protein | #N/A | #N/A | Smlt4383 | 98.7 | nuclease |
| Putative T4SS effector | H87-40_03005 | hypothetical protein | #N/A | #N/A | Smlt0113 | 99.4 | - |
| Putative T4SS effector | H87-40_03086 | hypothetical protein | #N/A | #N/A | Smlt0193 | 99.1 | lysozyme-like hydrolase |
| Putative T4SS effector | H87-40_03339 | hypothetical protein | #N/A | #N/A | Smlt4383 | 28.6 | nuclease |
| Putative T4SS effector | H87-40_03341 | hypothetical protein | #N/A | #N/A | Smlt0502 | 98.4 | - |
| Putative T4SS effector | H87-40_03343 | hypothetical protein | #N/A | #N/A | Smlt0502 | 33.5 | - |
| Putative T4SS effector | H87-40_03346 | hypothetical protein | #N/A | #N/A | Smlt0505 | 97.7 | - |
| T2SS | H87-40_01123 | Type II secretion system protein F | general secretion pathway protein F | K02455 | - | - | - |
| T2SS | H87-40-1b_01124 | putative type II secretion system protein HxcR | general secretion pathway protein E [EC:7.4.2.8] | K02454 | - | - | - |
| T2SS | H87-40_01129 | Type II secretion system protein G | general secretion pathway protein G | K02456 | - | - | - |
| T2SS | H87-40_02052 | Type II secretion system protein F | type IV pilus assembly protein PilC | K02653 | - | - | - |
| T2SS | H87-40_03507 | Type II secretion system protein E | general secretion pathway protein E [EC:7.4.2.8] | K02454 | - | - | - |
| T2SS | H87-40_03508 | Type II secretion system protein F | general secretion pathway protein F | K02455 | - | - | - |
| T2SS | H87-40_03509 | Type II secretion system protein G | general secretion pathway protein G | K02456 | - | - | - |

| Category | Gene locus | Gene annotation (Prokka) | Gene annotation (KEGG) | KEGG Orthology ID | Reference gene locus* | Protein sequence identity between H87-40-b and Reference gene locus* | Additional Information† |
|---|---|---|---|---|---|---|---|
| T4SS | H87-6_00915 | hypothetical protein | type IV secretion system protein VirD4 [EC:7.4.2.8] | K03205 | - | - | - |
| T4SS | H87-6_01399 | hypothetical protein | type IV secretion system protein VirD4 [EC:7.4.2.8] | K03205 | - | - | - |
| T4SS | H87-33_00775 | Type IV secretion system protein virB4 | type IV secretion system protein TrbE [EC:7.4.2.8] | K20530 | - | - | - |
| T4SS | H87-33_01947 | Type IV secretion system protein VirB11 | type IV secretion system protein VirB11 [EC:7.4.2.8] | K03196 | - | - | - |
| T4SS | H87-33_01948 | Type IV secretion system protein virB10 | type IV secretion system protein VirB10 | K03195 | - | - | - |
| T4SS | H87-33_01950 | Type IV secretion system protein virB8 | type IV secretion system protein VirB8 | K03203 | - | - | - |
| T4SS | H87-33_01953 | Type IV secretion system protein virB5 | type IV secretion system protein VirB5 | K03200 | - | - | - |
| T4SS | H87-33_01959 | Type IV secretion system protein virB4 | type IV secretion system protein VirB4 [EC:7.4.2.8] | K03199 | - | - | - |
| T2SS | H87-33_02211 | Type II secretion system protein G | Type II secretion system protein G | K02456 | - | - | - |
| T2SS | H87-33_02214 | Type II secretion system protein J | Type II secretion system protein J | K02459 | - | - | - |
| T2SS | H87-33_02221 | Type II secretion system protein F | Type II secretion system protein F | K02455 | - | - | - |

* Smlt: S. maltophila K279a; ICJ04: Stenotrophomonas sp. 169

†Bayer-Santos et al. (2019) PLos Pathogens https://doi.org/10.1371/journal.ppat.1007651

Supplementary Table 6. Endophytic bacterial community of plants grown in soil from which the selected strains were isolated.

| Taxonomy | Relative abundance |
|---|---|
| *Actinobacteriota;Actinobacteria;Micromonosporales;Micromonosporaceae;Other* | 27.4% |
| *Proteobacteria;Gammaproteobacteria;Burkholderiales;Comamonadaceae;Other* | 22.0% |
| *Chloroflexi;Ktedonobacteria;Ktedonobacterales;Ktedonobacteraceae;Other* | 4.0% |
| *Proteobacteria;Gammaproteobacteria;Burkholderiales;Comamonadaceae;Azohydromonas* | 3.6% |
| *Proteobacteria;Gammaproteobacteria;Xanthomonadales;Xanthomonadaceae;Pseudoxanthomonas* | 2.9% |
| *Actinobacteriota;Actinobacteria;Streptomycetales;Streptomycetaceae;Streptomyces* | 2.2% |
| *Proteobacteria;Gammaproteobacteria;Xanthomonadales;Rhodanobacteraceae;Tahibacter* | 2.0% |
| *Proteobacteria;Gammaproteobacteria;Burkholderiales;Oxalobacteraceae;Other* | 1.9% |
| *Proteobacteria;Alphaproteobacteria;Dongiales;Dongiaceae;Dongia* | 1.9% |
| *Proteobacteria;Gammaproteobacteria;Burkholderiales;Burkholderiaceae;Burkholderia-Caballeronia-Paraburkholderia* | 1.8% |
| *Proteobacteria;Alphaproteobacteria;Rhizobiales;Rhizobiaceae;Allorhizobium-Neorhizobium-Pararhizobium-Rhizobium* | 1.7% |
| *Proteobacteria;Alphaproteobacteria;Azospirillales;Azospirillaceae;Azospirillum* | 1.6% |
| *Bacteroidota;Bacteroidia;Cytophagales;Microscillaceae;Other* | 1.4% |
| *Proteobacteria;Gammaproteobacteria;Gammaproteobacteria Incertae Sedis;Unknown Family;Acidibacter* | 1.4% |
| *Bacteroidota;Bacteroidia;Chitinophagales;Chitinophagaceae;Niastella* | 1.1% |
| *Verrucomicrobiota;Verrucomicrobiae;Pedosphaerales;Pedosphaeraceae;uncultured* | 1.0% |
| *Proteobacteria;Alphaproteobacteria;Rhizobiales;Xanthobacteraceae;Bradyrhizobium* | 1.0% |
| *Proteobacteria;Alphaproteobacteria;Rhizobiales;Beijerinckiaceae;Methylobacterium-Methylorubrum* | 0.81% |
| *Proteobacteria;Alphaproteobacteria;Sphingomonadales;Sphingomonadaceae;Sphingomonas* | 0.70% |
| *Proteobacteria;Alphaproteobacteria;Micavibrionales;Other;Other* | 0.64% |
| *Proteobacteria;Gammaproteobacteria;Burkholderiales;Oxalobacteraceae;Massilia* | 0.60% |
| *Proteobacteria;Alphaproteobacteria;Caulobacterales;Caulobacteraceae;Asticcacaulis* | 0.54% |
| *Proteobacteria;Gammaproteobacteria;Xanthomonadales;Rhodanobacteraceae;Dyella* | 0.52% |
| *Patescibacteria;Saccharimonadia;Saccharimonadales;Other;Other* | 0.52% |
| *Bacteroidota;Bacteroidia;Chitinophagales;Chitinophagaceae;Other* | 0.51% |
| *Proteobacteria;Alphaproteobacteria;Rhizobiales;Rhizobiaceae;Other* | 0.50% |
| *Proteobacteria;Alphaproteobacteria;Sphingomonadales;Sphingomonadaceae;Novosphingobium* | 0.46% |
| *Bacteroidota;Bacteroidia;Cytophagales;Microscillaceae;uncultured* | 0.45% |
| *Proteobacteria;Gammaproteobacteria;Pseudomonadales;Pseudomonadaceae;Pseudomonas* | 0.37% |
| *Bdellovibrionota;Bdellovibrionia;Bdellovibrionales;Bdellovibrionaceae;Bdellovibrio* | 0.36% |
| *Bacteroidota;Bacteroidia;Chitinophagales;Chitinophagaceae;Flavisolibacter* | 0.35% |
| *Proteobacteria;Gammaproteobacteria;Burkholderiales;Rhodocyclaceae;Other* | 0.34% |
| *Actinobacteriota;Actinobacteria;Micrococcales;Microbacteriaceae;Leifsonia* | 0.33% |
| *Proteobacteria;Alphaproteobacteria;Sphingomonadales;Sphingomonadaceae;Other* | 0.33% |
| *Planctomycetota;Planctomycetes;Gemmatales;Gemmataceae;Gemmata* | 0.33% |
| *Cyanobacteria;Vampirivibrionia;Obscuribacterales;Obscuribacteraceae;Other* | 0.32% |
| *Chloroflexi;Anaerolineae;SBR1031;A4b;Other* | 0.31% |
| *Myxococcota;Polyangia;Haliangiales;Haliangiaceae;Haliangium* | 0.31% |
| *Verrucomicrobiota;Verrucomicrobiae;Chthoniobacterales;Chthoniobacteraceae;Chthoniobacter* | 0.30% |
| *Bacteroidota;Bacteroidia;Sphingobacteriales;env.OPS 17;Other* | 0.29% |
| *Proteobacteria;Alphaproteobacteria;Sphingomonadales;Sphingomonadaceae;Sphingopyxis* | 0.27% |
| *Proteobacteria;Gammaproteobacteria;Burkholderiales;Methylophilaceae;MM2* | 0.27% |
| *Verrucomicrobiota;Verrucomicrobiae;Opitutales;Opitutaceae;Lacunisphaera* | 0.26% |
| *Proteobacteria;Gammaproteobacteria;Burkholderiales;Oxalobacteraceae;Noviherbaspirillum* | 0.26% |
| *Proteobacteria;Alphaproteobacteria;Rhizobiales;Hyphomicrobiaceae;Hyphomicrobium* | 0.25% |
| *Proteobacteria;Alphaproteobacteria;Caulobacterales;Caulobacteraceae;Caulobacter* | 0.24% |
| *Proteobacteria;Gammaproteobacteria;Xanthomonadales;Rhodanobacteraceae;Other* | 0.24% |
| *Proteobacteria;Alphaproteobacteria;Rhizobiales;Xanthobacteraceae;Other* | 0.23% |
| *Proteobacteria;Alphaproteobacteria;Rhizobiales;Devosiaceae;Other* | 0.23% |
| *Planctomycetota;Planctomycetes;Planctomycetales;uncultured;Other* | 0.22% |
| *Proteobacteria;Alphaproteobacteria;Micropepsales;Micropepsaceae;uncultured* | 0.21% |
| *Bacteroidota;Bacteroidia;Cytophagales;Cytophagaceae;Sporocytophaga* | 0.20% |
| *Proteobacteria;Alphaproteobacteria;Rhizobiales;Xanthobacteraceae;Pseudolabrys* | 0.20% |
| *Bacteroidota;Bacteroidia;Cytophagales;Microscillaceae;Ohtaekwangia* | 0.19% |
| *Actinobacteriota;Actinobacteria;Micromonosporales;Micromonosporaceae;Actinoplanes* | 0.18% |
| *Actinobacteriota;Actinobacteria;Micrococcales;Microbacteriaceae;Other* | 0.18% |
| *Verrucomicrobiota;Verrucomicrobiae;Opitutales;Opitutaceae;Opitutus* | 0.18% |
| *Actinobacteriota;Actinobacteria;Propionibacteriales;Nocardioidaceae;Nocardioides* | 0.18% |
| *Cyanobacteria;Vampirivibrionia;Vampirovibrionales;Vampirovibrionaceae;Other* | 0.18% |
| *Planctomycetota;Planctomycetes;Planctomycetales;Rubinisphaeraceae;SH-PL14* | 0.17% |
| *Planctomycetota;Phycisphaerae;Tepidisphaerales;WD2101 soil group;Other* | 0.17% |

| Taxonomy | Relative abundance |
|---|---|
| *Proteobacteria;Alphaproteobacteria;Ferrovibrionales;Ferrovibrionaceae;Ferrovibrio* | 0.16% |
| *Proteobacteria;Alphaproteobacteria;Micropepsales;Micropepsaceae;Other* | 0.16% |
| *Actinobacteriota;Thermoleophilia;Solirubrobacterales;67-14;Other* | 0.14% |
| *Proteobacteria;Alphaproteobacteria;Sphingomonadales;Sphingomonadaceae;Sphingobium* | 0.14% |
| *Verrucomicrobiota;Verrucomicrobiae;Opitutales;Opitutaceae;Other* | 0.14% |
| *Proteobacteria;Alphaproteobacteria;Rhizobiales;Beijerinckiaceae;Bosea* | 0.14% |
| *Proteobacteria;Gammaproteobacteria;Xanthomonadales;Rhodanobacteraceae;Dokdonella* | 0.13% |
| *Proteobacteria;Alphaproteobacteria;Rhodospirillales;Magnetospirillaceae;Magnetospirillum* | 0.12% |
| *Proteobacteria;Gammaproteobacteria;Burkholderiales;Comamonadaceae;Hydrogenophaga* | 0.12% |
| *Proteobacteria;Gammaproteobacteria;Other;Other;Other* | 0.12% |
| *Bacteroidota;Bacteroidia;Flavobacteriales;Flavobacteriaceae;Flavobacterium* | 0.10% |
| *Armatimonadota;Armatimonadia;Armatimonadales;Other;Other* | 0.10% |
| *Myxococcota;Polyangia;Polyangiales;BIrii41;Other* | 0.10% |
| *Proteobacteria;Gammaproteobacteria;Gammaproteobacteria Incertae Sedis;Unknown Family;Candidatus Ovatusbacter* | 0.10% |
| *Bacteroidota;Bacteroidia;Cytophagales;Other;Other* | 0.10% |
| *Actinobacteriota;Thermoleophilia;Solirubrobacterales;Solirubrobacteraceae;Conexibacter* | 0.092% |
| *Proteobacteria;Gammaproteobacteria;Steroidobacterales;Steroidobacteraceae;Other* | 0.092% |
| *Proteobacteria;Other;Other;Other;Other* | 0.086% |
| *Armatimonadota;Fimbriimonadia;Fimbriimonadales;Fimbriimonadaceae;Other* | 0.086% |
| *Bacteroidota;Bacteroidia;Chitinophagales;Chitinophagaceae;Chitinophaga* | 0.078% |
| *Proteobacteria;Alphaproteobacteria;Caulobacterales;Caulobacteraceae;Other* | 0.076% |
| *Proteobacteria;Gammaproteobacteria;Xanthomonadales;Xanthomonadaceae;Luteimonas* | 0.074% |
| *Planctomycetota;Planctomycetes;Gemmatales;Gemmataceae;Fimbriiglobus* | 0.073% |
| *Myxococcota;Polyangia;Polyangiales;Polyangiaceae;Other* | 0.070% |
| *Bacteroidota;Bacteroidia;Chitinophagales;Chitinophagaceae;Terrimonas* | 0.069% |
| *Myxococcota;Polyangia;Polyangiales;Sandaracinaceae;uncultured* | 0.067% |
| *Planctomycetota;Planctomycetes;Pirellulales;Pirellulaceae;Pirellula* | 0.067% |
| *Proteobacteria;Alphaproteobacteria;Rhizobiales;Rhizobiaceae;Mesorhizobium* | 0.066% |
| *Planctomycetota;Phycisphaerae;Tepidisphaerales;CPla-3 termite group;Other* | 0.065% |
| *Bacteroidota;Bacteroidia;Chitinophagales;Chitinophagaceae;Pseudoflavitalea* | 0.064% |
| *Myxococcota;Myxococcia;Myxococcales;Myxococcaceae;P3OB-42* | 0.062% |
| *Proteobacteria;Gammaproteobacteria;Xanthomonadales;Rhodanobacteraceae;Luteibacter* | 0.062% |
| *Proteobacteria;Alphaproteobacteria;Caulobacterales;Caulobacteraceae;Phenylobacterium* | 0.060% |
| *Actinobacteriota;Actinobacteria;Micrococcales;Microbacteriaceae;Gryllotalpicola* | 0.058% |
| *Proteobacteria;Gammaproteobacteria;Xanthomonadales;Xanthomonadaceae;Other* | 0.057% |
| *Verrucomicrobiota;Verrucomicrobiae;Pedosphaerales;Pedosphaeraceae;Other* | 0.049% |
| *Proteobacteria;Alphaproteobacteria;Rhodospirillales;uncultured;Other* | 0.048% |
| *Verrucomicrobiota;Verrucomicrobiae;Pedosphaerales;Pedosphaeraceae;Ellin516* | 0.045% |
| *Proteobacteria;Gammaproteobacteria;Burkholderiales;Methylophilaceae;Other* | 0.043% |
| *Proteobacteria;Gammaproteobacteria;Xanthomonadales;Xanthomonadaceae;Lysobacter* | 0.042% |
| *Actinobacteriota;Actinobacteria;Catenulisporales;Actinospicaceae;Actinospica* | 0.041% |
| *Planctomycetota;Planctomycetes;Pirellulales;Pirellulaceae;uncultured* | 0.040% |
| *Acidobacteriota;Vicinamibacteria;Vicinamibacterales;uncultured;Other* | 0.038% |
| *Proteobacteria;Alphaproteobacteria;Rhizobiales;Xanthobacteraceae;uncultured* | 0.037% |
| *Verrucomicrobiota;Verrucomicrobiae;Chthoniobacterales;Terrimicrobiaceae;Terrimicrobium* | 0.037% |
| *Firmicutes;Bacilli;Bacillales;Bacillaceae;Bacillus* | 0.037% |
| *Planctomycetota;Phycisphaerae;Tepidisphaerales;Tepidisphaeraceae;Other* | 0.037% |
| *Verrucomicrobiota;Verrucomicrobiae;Chthoniobacterales;Chthoniobacteraceae;Other* | 0.036% |
| *Firmicutes;Clostridia;Clostridiales;Clostridiaceae;Clostridium sensu stricto 1* | 0.035% |
| *Bacteroidota;Bacteroidia;Cytophagales;Cytophagaceae;Other* | 0.033% |
| *Bacteroidota;Bacteroidia;Flavobacteriales;Weeksellaceae;Chryseobacterium* | 0.033% |
| *Proteobacteria;Alphaproteobacteria;Caulobacterales;Caulobacteraceae;Brevundimonas* | 0.032% |
| *Actinobacteriota;Actinobacteria;Corynebacteriales;Mycobacteriaceae;Mycobacterium* | 0.030% |
| *Bacteroidota;Bacteroidia;Chitinophagales;Chitinophagaceae;Ferruginibacter* | 0.029% |
| *Bacteroidota;Bacteroidia;Flavobacteriales;Crocinitomicaceae;Fluviicola* | 0.029% |
| *Chloroflexi;TK10;Other;Other;Other* | 0.028% |
| *Proteobacteria;Gammaproteobacteria;Burkholderiales;Alcaligenaceae;Other* | 0.028% |
| *Proteobacteria;Gammaproteobacteria;Steroidobacterales;Steroidobacteraceae;Steroidobacter* | 0.027% |
| *Actinobacteriota;Acidimicrobiia;Microtrichales;Iamiaceae;Iamia* | 0.027% |
| *Bdellovibrionota;Oligoflexia;0319-6G20;Other;Other* | 0.027% |
| *Proteobacteria;Gammaproteobacteria;Enterobacterales;Enterobacteriaceae;Other* | 0.025% |
| *Acidobacteriota;Acidobacteriae;Acidobacteriales;Acidobacteriaceae (Subgroup 1);Edaphobacter* | 0.025% |
| *Proteobacteria;Alphaproteobacteria;Rhizobiales;Beijerinckiaceae;Other* | 0.025% |
| *Proteobacteria;Gammaproteobacteria;Burkholderiales;Comamonadaceae;Variovorax* | 0.024% |

| Taxonomy | Relative abundance |
|---|---|
| *Verrucomicrobiota;Chlamydiae;Chlamydiales;cvE6;Other* | 0.023% |
| *Proteobacteria;Gammaproteobacteria;Burkholderiales;Rhodocyclaceae;Uliginosibacterium* | 0.022% |
| *Desulfobacterota;Desulfuromonadia;Geobacterales;Geobacteraceae;Citrifermentans* | 0.022% |
| *Proteobacteria;Alphaproteobacteria;Other;Other;Other* | 0.021% |
| *Proteobacteria;Gammaproteobacteria;Burkholderiales;Rhodocyclaceae;Candidatus Accumulibacter* | 0.021% |
| *Bacteroidota;Bacteroidia;Chitinophagales;Chitinophagaceae;Flavitalea* | 0.021% |
| *Bacteroidota;Bacteroidia;Cytophagales;Microscillaceae;Chryseolinea* | 0.020% |
| *Planctomycetota;Planctomycetes;Planctomycetales;Schlesneriaceae;Planctopirus* | 0.020% |
| *Planctomycetota;Planctomycetes;Pirellulales;Pirellulaceae;Pir4 lineage* | 0.019% |
| *Proteobacteria;Alphaproteobacteria;Rhizobiales;Pleomorphomonadaceae;Pleomorphomonas* | 0.018% |
| *Proteobacteria;Alphaproteobacteria;Sphingomonadales;Sphingomonadaceae;Blastomonas* | 0.017% |
| *Proteobacteria;Alphaproteobacteria;Rhizobiales;Rhizobiaceae;Ensifer* | 0.016% |
| *Myxococcota;Polyangia;Polyangiales;Sandaracinaceae;Sandaracinus* | 0.016% |
| *Acidobacteriota;Holophagae;Holophagales;Holophagaceae;Other* | 0.016% |
| *Planctomycetota;Phycisphaerae;Tepidisphaerales;Other;Other* | 0.016% |
| *Planctomycetota;Planctomycetes;Isosphaerales;Isosphaeraceae;Paludisphaera* | 0.015% |
| *Dependentiae;Babeliae;Babeliales;Other;Other* | 0.015% |
| *Proteobacteria;Gammaproteobacteria;Burkholderiales;Rhodocyclaceae;Azospira* | 0.015% |
| *Bacteroidota;Bacteroidia;Chitinophagales;Chitinophagaceae;Edaphobaculum* | 0.015% |
| *Proteobacteria;Alphaproteobacteria;Caulobacterales;Hyphomonadaceae;Hirschia* | 0.014% |
| *Proteobacteria;Gammaproteobacteria;Burkholderiales;Chromobacteriaceae;Other* | 0.014% |
| *Proteobacteria;Gammaproteobacteria;Burkholderiales;Oxalobacteraceae;Duganella* | 0.013% |
| *Proteobacteria;Gammaproteobacteria;Burkholderiales;Rhodocyclaceae;Dechloromonas* | 0.013% |
| *Chloroflexi;Anaerolineae;SBR1031;Other;Other* | 0.010% |
| *Proteobacteria;Alphaproteobacteria;Caulobacterales;Hyphomonadaceae;SWB02* | 0.010% |
| *Proteobacteria;Gammaproteobacteria;Pseudomonadales;Moraxellaceae;Acinetobacter* | 0.009% |
| *Spirochaetota;Leptospirae;Leptospirales;Leptospiraceae;Leptospira* | 0.009% |
| *Proteobacteria;Gammaproteobacteria;Burkholderiales;Chromobacteriaceae;Vogesella* | 0.008% |
| *Proteobacteria;Gammaproteobacteria;Methylococcales;Methylomonadaceae;Methylomonas* | 0.007% |
| *Bacteroidota;Bacteroidia;Bacteroidales;Prolixibacteraceae;WCHB1-32* | 0.007% |
| *Firmicutes;Bacilli;Paenibacillales;Paenibacillaceae;Paenibacillus* | 0.006% |
| *Proteobacteria;Gammaproteobacteria;Burkholderiales;Burkholderiaceae;Cupriavidus* | 0.005% |
| *Bdellovibrionota;Bdellovibrionia;Bacteriovoracales;Bacteriovoracaceae;Peredibacter* | 0.004% |
| *Proteobacteria;Alphaproteobacteria;Rhizobiales;Rhizobiaceae;Brucella* | 0.004% |
| *Verrucomicrobiota;Verrucomicrobiae;Pedosphaerales;Pedosphaeraceae;DEV008* | 0.004% |
| *Acidobacteriota;Acidobacteriae;Paludibaculum;Other;Other* | 0.004% |
| *Actinobacteriota;Actinobacteria;Micrococcales;Micrococcaceae;Other* | 0.004% |
| *Campilobacterota;Campylobacteria;Campylobacterales;Sulfurospirillaceae;Sulfurospirillum* | 0.004% |
| *Myxococcota;Myxococcia;Myxococcales;Anaeromyxobacteraceae;Anaeromyxobacter* | 0.004% |
| *Desulfobacterota;Desulfovibrionia;Desulfovibrionales;Desulfovibrionaceae;Desulfovibrio* | 0.003% |
| *Bacteroidota;Bacteroidia;Bacteroidales;Other;Other* | 0.003% |

Taxonomic information was obtained by 16S rRNA amplicon sequencing.

Supplementary Table 7. Five types of media were used to isolate plant-associated microbial complexes.

| Medium | Reagent | Amount (per 1L) |
|---|---|---|
| R2A broth | R2A broth “DAIGO” (Nihon Pharm. Co., Japan) | 3.2 g |
| | Agar | 15 g |
| NYGB (Nutrient yeast extract glycerol broth) | Peptone | 5 g |
| | Yeast extract | 3 g |
| | Glycerol | 20 mL |
| | Agar | 15 g |
| | (pH 7) | |
| TSB (Typtic soy broth) | Casein peptone | 17 g |
| | Soy peptone | 3 g |
| | NaCl | 5 g |
| | $K_2HPO_4$ | 2.5 g |
| | D(+)-glucose | 2.5 g |
| | Agar | 15 g |
| | (pH 7) | |
| PDA (Potato dextrose agar) | Potato dextrose agar (Difco co., USA) | 24 g |
| WA (Water agar) | Agar | 15 g |

NYGB, TSB and PDA are nutrient-rich media, on the other hand, R2A and WA are nutrient-poor media.
